# LRRTM4 controls cell migration via Glypican interactions

**DOI:** 10.64898/2026.04.14.718450

**Authors:** Claudia Peregrina, Miguel Berbeira-Santana, Maria Carrasquero-Ordaz, Irene Rodríguez-Navarro, Kamel El Omari, Hao-Yi Li, Gönül Seyit-Bremer, Rüdiger Klein, Valerie Castellani, Elena Seiradake, Daniel del Toro

**Affiliations:** Department of Biomedical Sciences, Faculty of Medicine and Health Sciences, Institute of Neurosciences, IDIBAPS, CIBERNED, University of Barcelona, Barcelona, Spain; Department of Biochemistry, University of Oxford, Oxford, OX1 3QU, UK; Kavli Institute for Nanoscience Discovery, University of Oxford, Oxford, OX1 3QU, UK; Diamond Light Source, Harwell Science and Innovation Campus, Didcot OX11 0DE, UK; Institute of Precision Medicine, College of Medicine, National Sun Yat-Sen University, Kaohsiung, Taiwan; Department of Molecules-Signaling-Development, Max-Planck Institute for Biological Intelligence, 82152 Martinsried, Germany; MeLis, University of Lyon, Université Claude Bernard Lyon 1, CNRS UMR 5284, INSERM U1314, Institut NeuroMyoGène, 8 avenue Rockefeller 69008 LYON, France

**Keywords:** LRRTM, glypicans, GPC4, GPC3, LRRTM4, cell guidance, cortex development, cancer migration

## Abstract

Leucine-rich-repeat-transmembrane neuronal proteins (LRRTMs) play a major role in neuronal connectivity and synapse formation, but their function during early development remains largely unexplored and roles in cell migration have not been described. Here, we identify LRRTM4 as a major coordinator of cell migration within the developing cortex and in cancer cells. Using structural predictions and site-directed mutagenesis to abolish the LRRTM4-glypican interaction, we show that LRRTMs interact with the core domain of glypicans, which, in the midgestational murine cortex, are presented by radial glia cells. We demonstrate that glypican-bound LRRTM4 mediates contact-repulsion in migrating neurons and cancer cells *in vitro*. Our results demonstrate that LRRTMs are used by migrating cells responding to glypicans, and that this signaling system is active in migrating cancer cells and during development.

## Introduction

Cell migration is a fundamental process that shapes tissue organization during development and is frequently hijacked in pathological contexts such as cancer metastasis^1^. In the developing cerebral cortex, newly generated neurons migrate along radial glial scaffolds to reach their final laminar positions, a process controlled by a complex combination of extracellular cues, receptor complexes and cell-cell interactions^2–5^. Interestingly, many of the molecular mechanisms that regulate neuronal migration are shared with those driving tumor cell motility and invasion, including receptors involved in neuron-neuron and neuron-glia interactions^6–8^. These cell-cell communications rely on cell adhesion molecules and extracellular guidance signals that direct cell movement through attractive or repulsive interactions^5,9^. In the brain, such interactions are critical not only during development but also for neuron connectivity and synaptic activity in the adult brain^10^. Indeed, we and others have shown that several proteins initially identified as synaptic adhesion molecules play a role at earlier time points, much before synapse development, including neuronal migration and axon guidance ^11–13^. In this context, combinatorial interactions between receptors is a key feature, as relatively few neuronal guidance receptors participate in multiple processes, including neuronal migration, axon guidance and synapse formation^5,14^. Moreover, the cellular responses elicited by these interactions are context dependent, such that receptors that promote adhesion during synapse development can instead mediate repulsion in migrating cells or navigating axons ^12,15,16^.

One prominent class of cell surface receptors that exhibits such functional versatility are those containing extracellular Leucine-rich repeat (LRR) domains. Several LRR protein families have emerged as key regulators of tissue morphogenesis, participating in processes ranging from neuronal migration and axon guidance to circuit assembly and synapse formation^14,17–19^. A prime example comes from FLRT(1-3) receptors which bind multiple receptor partners to regulate all these processes in a context-dependent manner. FLRTs can mediate homophilic adhesion or heterophilic repulsion through the formation of distinct protein complexes with Unc5, Latrophilin, and Teneurin receptors^11–13^. Similar to FLRTs, Leucine-rich-repeat-transmembrane neuronal proteins (LRRTM1-4) participate in synapse formation and circuit wiring by interacting with partners such as neurexin^19,20^. LRRTMs are single transmembrane receptors with an LRR domain in the N-terminal extracellular domain, followed by a transmembrane region and a cytoplasmic tail containing a C-terminal PDZ-binding motif^21^. LRRTMs are highly expressed in the brain and are also present during brain development before synapse formation^22^, it remains unknown whether they participate in processes such as neuronal migration, as observed for FLRTs^12,23^. Interestingly, recent studies implicate LRRTMs in functions beyond the nervous system, including roles for LRRTM4 in cancer proliferation and metastasis^24^. However, the underlying molecular mechanisms remain unclear.

Glypicans are a family of heparan sulfate proteoglycans (GPC1-6) that are bound to the cell surface by a glycosylphosphatidylinositol (GPI) anchor. These extracellular receptors are broadly expressed across tissues and are frequently upregulated in tumors, where they participate in several processes including morphogen signaling, cell proliferation, and cell migration^25–27^. All glypicans consist of a structured N-terminal extracellular core domain followed by a C-terminal linker region of approximately 80 amino acids. Structural studies have shown that the core domain adopts an α-helical fold organized into N-, M-, and C-terminal lobes stabilized by conserved disulfide bonds and containing a furin-like cleavage site^28^. The C-terminal region carries heparan sulfate (HS) glycan attachment sites and a GPI anchor that tethers glypicans to the cell surface^29^. Most glypican interaction partners bind to their HS glycans, including Wnts, Frizzled and Hedgehog^30–32^. Interestingly, some FLRT binding partners, such as Unc5 receptors, interact with the core protein of GPC3 through a glycan-glycan interaction involving a C-mannosylated tryptophan in Unc5D and an N-linked glycan on GPC3^8^. During brain development GPC3 is expressed by radial glia (RG) cells and regulates the migration of Unc5D-expressing neurons through interactions along radial glial processes^8^. Interestingly, this same interaction also contributes to the migration of tumor cells, such as neuroblastoma^8^. Interestingly, the removal of this interaction due to mutation of GPC3 resulted only in a partial phenotypic effect^8^ suggesting that additional receptors for GPCs are present in neurons and cancer cells^8^.

Here, we report that all LRRTMs bind to the core domain of glypicans, which are expressed in many types of migrating cells and their environments. The interaction site on GPC is distinct from the previously identified Unc5-binding site on GPC3^8^. In the developing cortex, glypicans are present in RG cells where they act as ligands for LRRTM4, which is expressed on migrating neurons. Using an AI-based structural prediction approach, we developed tools to manipulate this interaction and demonstrate that it controls cortical migration through a cell repulsion-mediated mechanism. Moreover, this interaction similarly regulates the migration of multiple cancer cell types. Together, our findings reveal shared migratory mechanisms that provide insights into both brain development and cancer progression.

## Results

### LRRTMs and Glypicans are expressed during cortical development

Extracellular proteins containing leucine-rich repeat (eLRR) domains are highly expressed in the adult nervous system, with specific expression patterns that often label distinct subpopulations of neurons^33^. We measured the abundance of eLRRs in the developing cortex at embryonic day 15.5 (E15.5, mid gestation) using a lectin-based pulldown followed by mass spectrometry (Fig. 1A), as previously described^13,34^. Among all LRR-containing receptors, we found that LRRTMs represent the second family, after FLRTs, with all members ranked within the top 25% of the most enriched receptors (Fig. 1B and S1A). To examine their expression pattern across cortical cell subpopulations, we analyzed our published single-cell RNA-seq (scRNA-seq) database obtained from dissociated cortical tissue at E15.5 (Fig. 1C). All LRRTM family members (LRRTM1-4) were preferentially expressed in neurons (Fig. 1D). Among them, LRRTM4 showed the highest enrichment in migrating neurons. Distribution analysis using categorized clusters showed that 65% of migrating neurons express LRRTM4, whereas fewer than 24% express any of the other LRRTM members (Fig. 1E). In the synaptic context, LRRTM proteins localize postsynaptically and bind *in trans* to presynaptic glypicans and neurexins^18^. We previously reported a similar trans-cellular configuration for other synaptic proteins, including FLRTs, Teneurins, Latrophilins in neuron-radial glia (RG) interactions during cortical development^12^. Based on this framework, we analyzed the expression of glypicans (GPC1-6) and neurexin (NRXN1) in cortical progenitor populations. We found that, similar to GPC3^8^, GPC4 is highly enriched in apical progenitors (APs) compared with intermediate progenitors (IPs), and relative to other glypicans such as GPC2, as well as NRXN1 (Fig. 1F-G). Consistent with the expression data, *in situ* hybridization (ISH) for GPC4 showed restricted expression within the germinal layers, predominantly in the ventricular zone (VZ), where AP cells are located at E15.5. In contrast, LRRTM4 showed strong enrichment in young/migrating neurons located in the intermediate zone (IZ) (Fig. 1H). Analysis of two independent scRNA-seq datasets^35,36^ further confirmed that LRRTM4 is enriched in migrating neurons, whereas AP cells show high expression of GPC4 from E13.5 to E17.5 (Fig. 1I, S1B-H). To corroborate the interaction between LRRTM4 and GPC4^20^, we used soluble LRRTM4 ectodomain as a bait in binding assays on transfected HeLa cells expressing GPC4 (Fig. 1J). Based on these results, we developed a working model in which migrating neurons expressing LRRTM4 interact with glypicans, such as GPC4, present in radial glia (RG) cells (Fig. 1K). We proceeded with *in vitro* functional analysis to understand the roles of these proteins in early cortical development.

**Figure 1.**
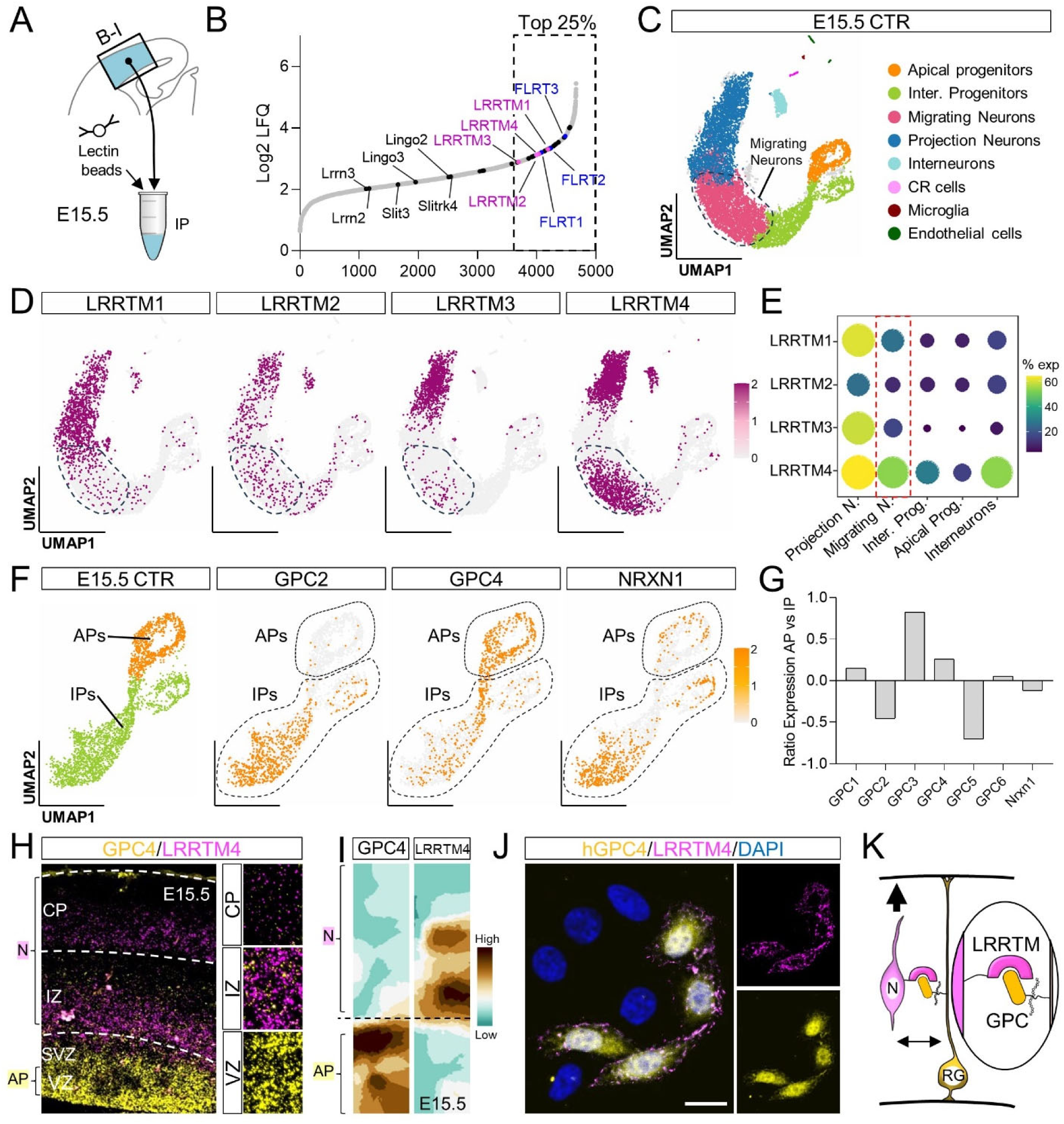
LRRTMs and glypicans are expressed during cortical development. (A) Schematic showing the cortical region used for lectin-based pulldown at E15.5. (B) Plot showing proteins captured by lectin pulldown and identified by mass spectrometry using label-free quantification (LFQ). The results of four separate sets of pull-downs were averaged. Dashed rectangle indicates the top 25% of the most enriched proteins. (C) UMAP derived from scRNA-seq analysis of cortical cells at E15.5 (GSE27179^73^), cells are colored by type. (D) UMAP visualization showing expression of LRRTM(1-4). (E) Dot plot quantifying the distribution of LRRTM1-4 positive cells across categorized clusters. (F) UMAP visualization showing expression of glypicans and neurexin (NRXN1) in the intermediate (IP) and apical progenitor (AP) cortical populations. (G) Quantification of expression ratio in APs vs IPs for glypicans and NRXN1. (H) Double in situ hybridization (ISH) for LRRTM4 (magenta) and GPC4 (yellow) shows their expression in the cortex at E15.5. The layers enriched in neurons (N) and apical progenitors (AP) are indicated. (I) mRNA levels obtained from single-cell transcriptome sequencing of embryonic cortical cells at E15.5. The Y axis indicates the differentiation state of individual cortical cells from an apical progenitor (AP) to a postmitotic neuron (N) deduced from transcriptome analysis^35^. (J) Immunofluorescence-based binding assay using HeLa cells transfected with hGPC4 (yellow) and incubated with exogenous LRRTM4 ectodomain (magenta). (K) Summary model showing LRRTM4 and GPC4 expression patterns. Scale bar 10um (J).

### LRRTM4-Glypican interaction produces contact repulsion *in vitro*

To study the effects of LRRTM4-GPC4 binding on cortical neuron migration, we performed time-lapse imaging of E15.5 embryonic cortical explants grown on GPC4-coated dishes and measured the migration of neurons exiting the explant (Fig. 2A). Using automatic tracking (Movie S1), we found that cortical neurons migrated slower and shorter distances on GPC4-coated surfaces compared with control surfaces (Fig. 2B-C). Given that LRRTM4 is expressed in migrating neurons and its binding partner GPC4 in RG cells, we next examined the functional role of this interaction in the context of neuron-RG fiber interactions. We used our previously described nanofiber assay^12^ (Fig. 2D), which consists of parallel, aligned nanofibers that mimic the fibrillary environment of RG fibers^37^. Cortical explants from E15.5 embryos, when placed onto nanofibers, displayed directed neuronal migration along the fibers (Fig. 2D, S2A and Movie S2). Using semi-automated analysis^12^, we found that neurons migrated shorter distances on GPC4-coated nanofibers (Fig. 2E-F). The reduced migration observed on GPC4-coated substrates may result from a contact-repulsion mechanism, similar to what we previously reported for another glypican family member, GPC3, which is also enriched in RG cells^8^. Using stripe assay, we found that GPC4 is strongly repulsive for cortical neurons (Fig. 2G-H). Addition of soluble LRRTM4 ectodomain to the medium significantly reduced the repulsion elicited by GPC4 (Fig. 2I-J). Notably, among all LRRTM members, LRRTM4 also induced strong repulsion of cortical cells (Fig. S2B-E), a behavior similar to that observed for FLRT3, a well-established regulator of cortical neuron migration^11,23^. Together, these results suggest that GPC4 induces repulsion of cortical neurons through an interaction with LRRTM4 (Fig. 3A).

**Figure 2.**
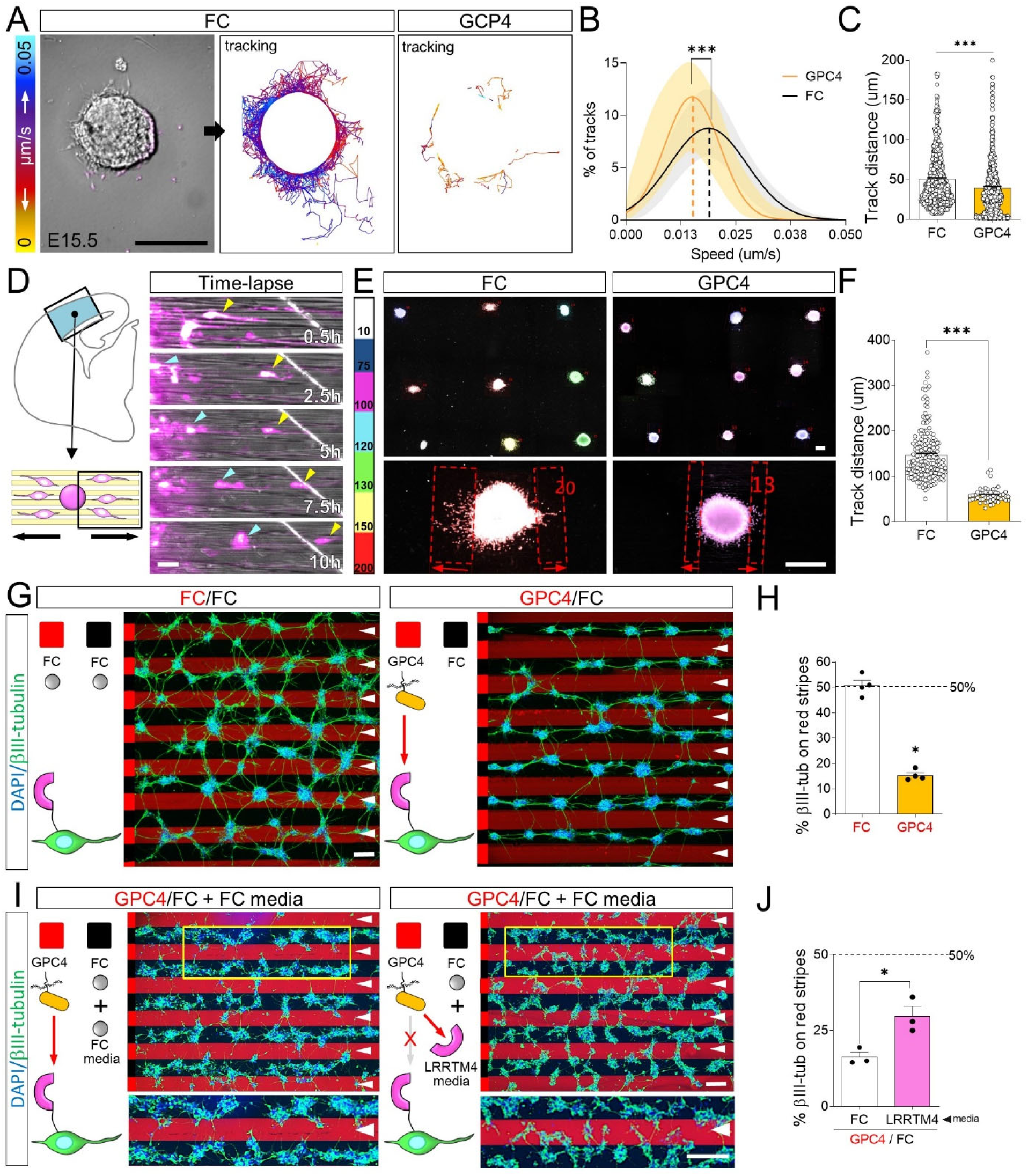
LRRTM4-GPC4 interaction regulates cell migration *in vitro* (A) Time-lapse analysis of cortical neurons exiting E15.5 cortical explants grown on surfaces coated with FC (control) or GPC4. An explant growing on FC coated surface is shown on the left. Neurons were tracked (lines) and color-coded based on migration speed (right). (B) Distribution of average migration speeds. n > 5 movies per condition. ***p<0.001, Student’s t test. (C) Quantification of total track distance from (A). ***p<0.001, Student’s t test. (D) Diagram depicting the nanofiber assay (left) and snapshots from a time-lapse video of GFP-expressing neurons (magenta) migrating on FC coated nanofibers. Cyan and Yellow arrowhead points to their cell soma (E) Explants grown on nanofibers coated with FC and GPC4 for 2 days in vitro (DIV). DAPI staining is color-coded based on the average distance from the explant, indicating migration length (see also Figure S2A). (F) Quantification of the data shown in (E). n >50 explants (3 experiments per condition). ∗∗∗p < 0.001, Student’s t test. (G) E15.5 dissociated cortical neurons grown on alternate stripes containing FC (black) and GPC4 proteins (red). Neurons were stained with anti-β-III-tubulin (green) and DAPI (nuclei, white). Red stripes are indicated by yellow arrowheads. (H) Quantification of the percentage of β-III-tubulin pixels on red stripes. n = 3 independent experiments. ∗p < 0.05, Student’s t test. (I) Similar to G but neurons were cultured with FC or LRRTM4 soluble proteins added to the medium. (J) Quantification of the data shown in (I). n = 3 different experiments. ∗p < 0.05, Student’s t test. Scale bar 300um (A, E), 20um (D), 100um (G,I).

**Figure 3.**
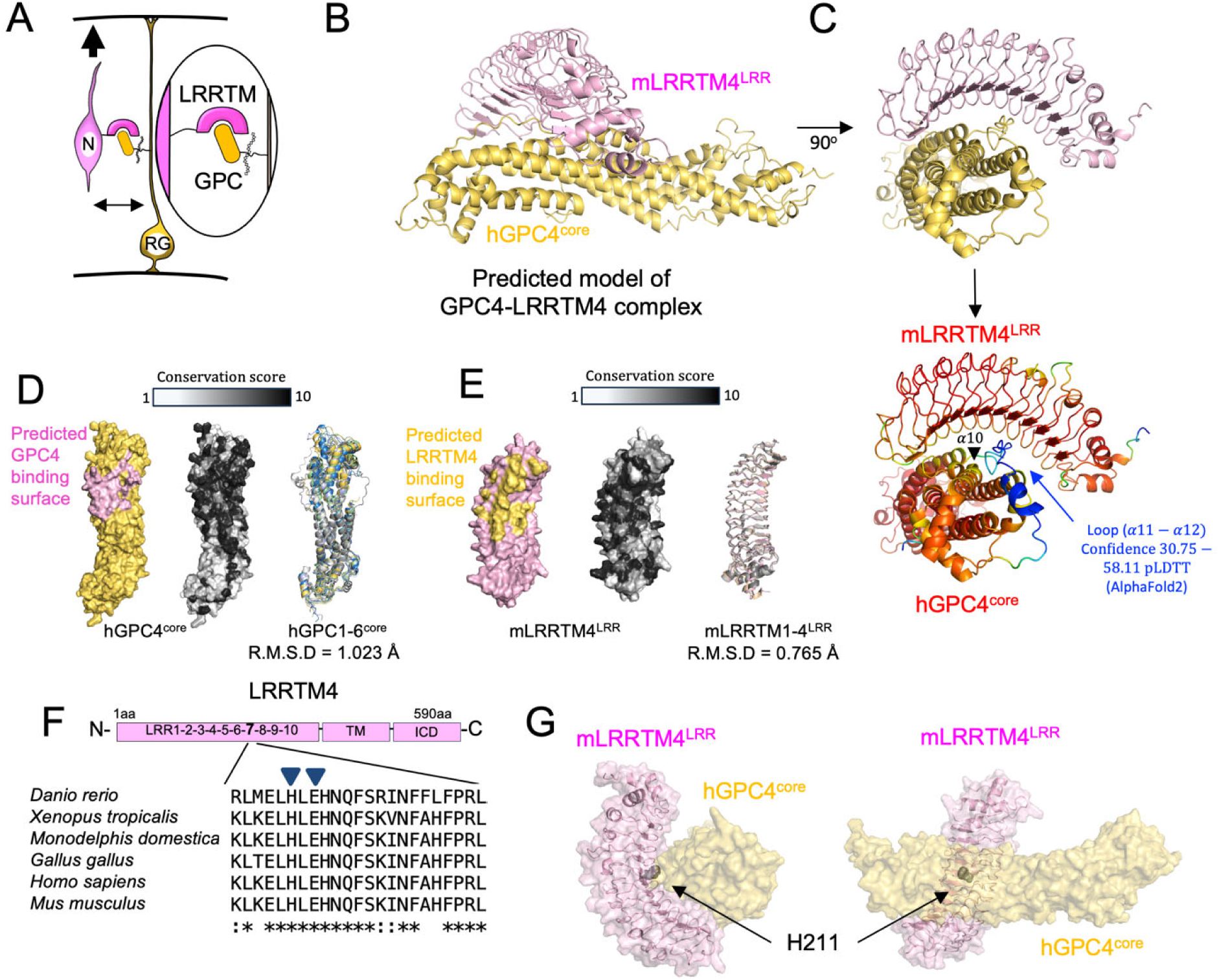
Predicted model of the GPC4^core^ – LRRTM4^LRR^ interaction and GPC4 mutagenesis. Structural analysis of the GPC4-LRRTM4 complex using AlphaFold. **(A)** A model depicting a migrating neuron (N) and a radial glia fiber (RG), expressing LRRTM4 and GPC4 proteins, respectively. We postulate an interaction between their extracellular domains based on data in Figures 1 and 2. **(B)** AlphaFold was used to predict structures of LRRTM4 LRR and GPC4 core, and their interaction mode. **(C)** As panel B, but the view is rotated by 90°. The model is coloured as in panel B (top) or according to the confidence scores calculated with AlphaFold (bottom). **(D)** GPC4 surface views and cartoon views are shown in the same orientation. Left: the predicted LRRTM binding surfaces are coloured in pink. Middle: the colours represent sequence conservation scores after aligning GPC4 sequences of *homo sapiens, mus musculus, gallus gallus, monodelphis domestica, xenopus tropicalis,* and *danio rerio*. Right: Predicted models of GPC1-6 are superposed. GPC1 (lime green), GPC2 (skyblue), GPC3 (grey), GPC4 (yellow), GPC5 (light grey), GPC6 (light blue). **(E)** Analogous to panel D, but showing LRRTM4 and homologues instead. LRRTM1 (light grey), LRRTM2 (wheat), LRRTM3 (grey), LRRTM4 (pink). **(F):** Multiple Sequence Alignment of the sequences used in panel E with the mutated residues highlighted. **(G)** The position of H211, where the nG mutation was introduced, is pointed out.

### Structural predictions of the LRRTM4 LRR and GPC4^core^ complex

The LRR domain is one of the most common protein domain repeats across species. It is a structural motif composed of tandem repeats of 20-30 amino acids that adopt a curved, conserved horseshoe-shaped structure. Many LRR-containing receptors bind their ligands through the concave surface of the LRR domain, being one of the most effective and evolutionary conserved protein-protein interaction motif^18,33^. Similarly, the three-dimensional structure of the glypican core protein is conserved^8,38–42^, despite only moderate amino acid sequence homology among family members, consistent with conserved roles in development and signaling across vertebrates. Thus, and given our failed attempts to solve the experimental structure due to difficulties with producing suitable protein samples, we applied AI-based structure prediction methods with AlphaFold to generate structural hypotheses for this interaction, an approach increasingly used to identify protein-protein interfaces^43–45^. After inputting sequence information (GPC4 residues K26-E536, LRRTM4 residues Q31-H423), we obtained a model that suggested a single major interface, in which the concave region of the LRRTM4 LRR domain predominantly contacts helix 10 and the extended loop located between helices 11 and 12 of GPC4 core (Fig. 3B-E). Note that the confidence score for this GPC4 loop region is lower than for the rest of the model (Fig. 3C). Interestingly, the predicted pose is different to the binding model for neurexin 1beta, which binds at the C-terminal end of the LRR domain of LRRTM2^21^. It is rather more similar to the binding modes of other LRR proteins, such as FLRT, which binds the Latrophilin receptor via the concave surface of the LRR domain^46^. The three-dimensional structures of GPC1-6 core domains, and LRRTM1-4 LRR domains are predicted to be structurally similar, respectively, and sequence alignments suggest that the predicted GPC4-LRRTM4 interaction surfaces are conserved in other GPCs and LRRTMs (Fig. 3D-F). Also, structural comparison shows that the interaction of GPC3 with Unc5^8^, which involves multiple distinct surface areas of GPC3, does not overlap with the predicted binding site for LRRTM (Fig. S3A,B). The main purpose of the structural analysis presented above was to underpin model-based protein engineering to disrupt the interaction for subsequent functional investigations *in vitro* and *in vivo*. We chose an established method, which introduces an N-linked glycosylation site at the binding interface^8,11,46–50^. We chose position 211 of LRRTM4 for the modification (H211N+E213T). We extended this mutagenesis to LRRTM2 as well (H211N+E213T). The site is located at the heart of the sequence-conserved interacting surface (Fig 3G), and was designed to impact the protein fold as little as possible. In analogy to this mutant, we also produced GPC4 mutants to introduce a glycosylation in position 104 or 318 (K104N+D106T, K318N+S320T) (Fig. S3C, D). We show that the mutants are expressed at the surface of cells at comparable levels to the wild type sequences (Fig. S4A, B), suggesting that they pass the strict quality control mechanisms that would inhibit misfolded protein from being secreted through the endoplasmatic reticulum and Golgi apparatus^51,52^. Going forwards, these mutants are referred to as GPC “nL” (for non-LRRTM-binding) and LRRTM “nG” (for non-GPC-binding).

### Cell-based aggregation assays confirm interactions via LRRTM and GPC proteins

We performed cell aggregation assays to investigate the binding properties of GPC4 and LRRTM constructs, including mutants, *in vitro*. In these assays, interacting cell surface receptors induce cell aggregation of normally suspended K562 cells (Fig. 4A). We verify cell surface presentation of our constructs using immunofluorescence (Fig. S4A, B). In agreement with previous reports showing that LRRTM4 binds several glypican family members^20^, and our sequence conservation analysis (Fig. 3D), the cell aggregation assays showed that LRRTM4-expressing cells adhere to cells expressing GPC3, GPC4, or GPC5, and that these interactions are abolished by the addition of exogenous GPC3 (Fig. 4A and S4B, E). We confirm this result using soluble proteins in a pulldown experiment, where GPC3^core^ effectively pulls down LRRTM2,4 and Unc5D proteins, but not the negative control protein, Latrophilin (Fig S4C). Together, these results support a model in which LRRTMs interacts with different glypicans through a conserved binding surface. This is further supported by cell aggregation assays, where the expression of wild type LRRTM2 or LRRTM4 promotes cell aggregation with GPC3-5 expressing cells, but not the non-GPC-binding “nG” mutants (Fig. 4B and S4E). Notably, the “nG” mutant does not affect cell aggregation mediated by LRRTM4 binding to Neurexin 1beta (Fig. 4C and S4), a well-characterized binding partner of LRRTM proteins^20^. Vice versa, the GPC4 “nL” mutant K104N+D106T showed poor cell adhesion to LRRTM1-4 (Fig. 4C-F and S4E). We confirmed that the same nL mutation, implemented into GPC3 (V109N + Q111T) removes LRRTM4-GPC3 mediated cell aggregation, further confirming the conserved nature of the interaction, while not affecting Unc5B binding (Fig. 4G and S4E). We conclude that LRRTM-GPC interactions can be controlled using specific mutations.

**Figure 4.**
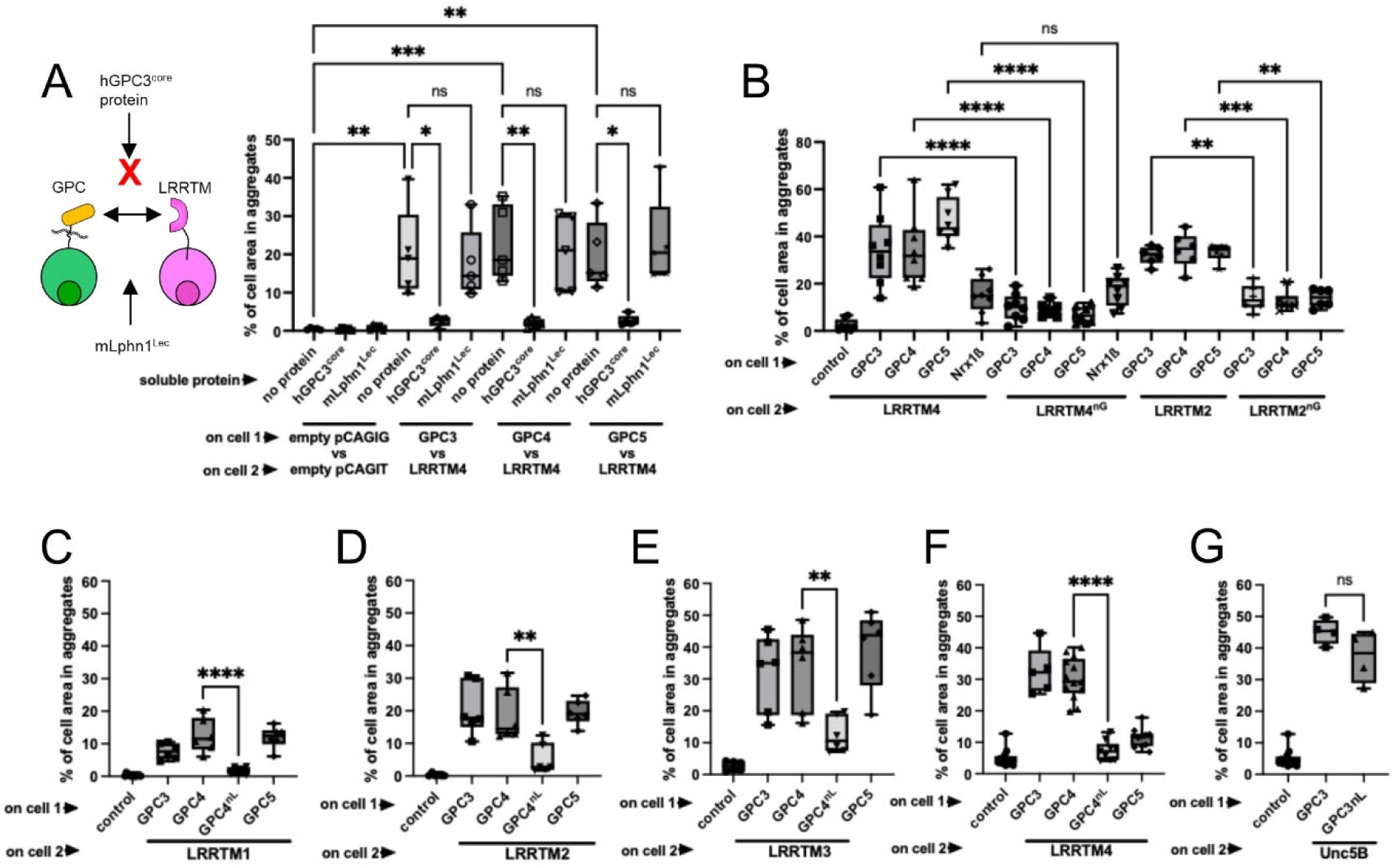
The GPC^core^ – LRRTM^LRR^ interaction is conserved. We used a cell-cell aggregation assay to assess LRRTM-GPC interactions *in trans*. In these experiments GPC-expressing cells aggregate with LRRTM-expressing cells if the two proteins interact. Representative images are shown in Figure S4. **(A)** The addition of soluble GCP3^core^ protein inhibits the GPC4-LRRTM4 interaction, presumably due to competition, while the addition of Latrophilin protein does not. **(B)** GPC and LRRTM homologues interact. The nG mutation in LRRTM2 or LRRTM4 abolishes effective cell aggregation. **(C)** The GPC nL mutation abolishes the cell aggregation between LRRTM1 and GPC4. **(D)** Same as panel C but for LRRTM2. **(E)** Same as panel C and D but for LRRTM3. **(F)** Same as panel C, D and E but for LRRTM4. **(G)** The GPC3 nL mutant does not abolish cell aggregation between Unc5B and GPC3. Note that the mutants still bind Unc5 and Neurexin, suggesting that the binding sites are distinct. n.s. = not significant. ****p < 0.0001. One-way ANOVA test with Tukey’s post hoc analysis (A,B,C,D,E,F,G). Box and whiskers plots (A,B,C,D,E,F,G) are defined as minimum data point to maximum data point, centre is median, percentiles are 25, 50, 75.

### LRRTM4-glypican interaction control cell migration in the developing cortex

Having established how to manipulate the LRRTM-GPC interaction, and that GPC4 is repulsive for LRRTM4-expressing migrating neurons *in vitro*, we next examined its role *in vivo*, using cortical development as a model system. During development, pyramidal neurons are born in the proliferative zone and radially migrate to settle in the cortical plate^53^. Using in utero electroporation (IUE), we knocked down endogenous LRRTM4 in E13.5 cortices by expressing small hairpin RNA (shRNA) target sequences into the pCAG-miR30 vector system^54^ (Fig. 5A-B). Analysis at E16.5 showed reduced migration, with fewer neurons reaching the upper layer of the cortical plate (Fig. 5C-D). Similar migration defects were observed following GPC4 knockdown (Fig. S5A-B and Fig. 5C-D), consistent with our previous findings showing that loss of GPC3 delays cortical migration^8^. When categorizing the neurons based on their morphologies (Fig. S5D-E), we did not observe significant differences between LRRTM4 and GPC4 knockdown neurons and tomato (TOM) controls. Overexpression of wild type LRRTM4 resulted in a more pronounced migration delay, with most neurons (∼60%) remaining in the intermediate zone and fewer entering the cortical plate (∼22%) compared with control conditions (∼73%) (Fig. 5E). Notably, introduction of the LRRTM4^nG^ mutant in these over-expression experiments fully rescued the inhibitory effect of LRRTM4 over-expression. These results indicate that glypicans are the primary binding partner mediating LRRTM4-dependent regulation of cortical migration, as the LRRTM4^nG^ mutant retains the ability to interact with neurexins (Fig. 4B-S4E).

**Figure 5.**
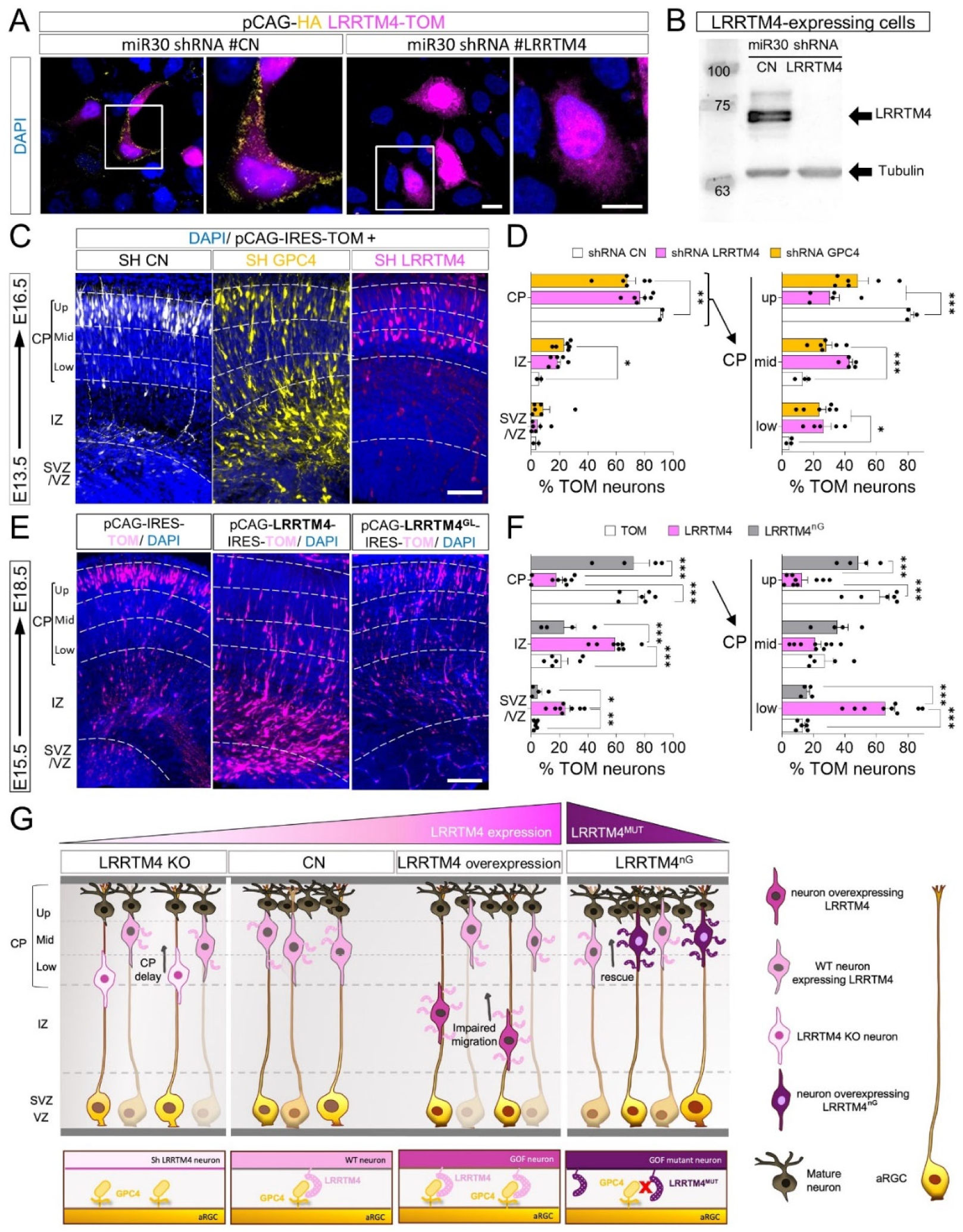
LRRTM4 interaction with GPC4 delays neuron migration. (A) HeLa cells were transfected with pCAG expressing HA-tagged LRRTM4 (magenta) together with pCAG-miR30 containing a control shRNA or an shRNA targeting murine LRRTM4. Cells were cultured for 2 DIV, and immunostained for surface LRRTM4 (Ha tag, yellow). A representative cell for each condition are shown, with magnified views on the right. (B) Anti-HA western blot showing expression levels of HA-tagged LRRTM4 in the same cells transfected in A. Tubulin was used as a loading control. (C) Coronal sections of an E16.5 cortex previously electroporated with pCAG-IRES-TOM and a pCAG-miR30 vector coding for shRNA control (CN), shRNA targeting murine GPC4 (yellow) or LRRTM4 (magenta). The cortical plate (CP) was defined based on DAPI staining. TOM-positive cells in the SVZ/VZ, IZ and CP were automatically identified and the percentage in each layer quantified. The CP was further subdivided into 3 bins (up, mid, low). (D) Quantification of data shown in (C). n = 3 shRNA CN, n = 6 shRNA LRRTM4, and n = 6 shRNA GPC4 electroporated brains. ∗p < 0.05, ∗∗p < 0.01, ∗∗∗p < 0.001, one-way ANOVA test with Tukey’s post hoc analysis. (E) Coronal sections of E16.5 cortex after IUE using empty vector (pCAGTOM, control), LRRTM4 and LRRTM4^LG^. TOM-positive cells were quantified for each bin. (F) Quantification of data shown in (E). n = 3 TOM, n = 8 LRRTM4, and n = 5 LRRTM4^nG^ electroporated brains. ∗p < 0.05, ∗∗p < 0.01, ∗∗∗p < 0.001, one-way ANOVA test with Tukey’s post hoc analysis. (G) Cartoon depicting how LRRTM4-Glypican interactions regulate cortical neuron migration. In the WT, radial migration is controlled by a balance between adhesive and repulsive interactions. LRRTM4 knockdown (KO) or overexpression disrupts this balance, resulting in impaired migration. In contrast, LRRTM4^nG^ overexpression produces a phenotype similar to WT condition, indicating that glypicans are the primary binding partner mediating LRRTM4-dependent regulation of cortical migration

We also over-expressed GPC4 using the same IUE approach (Fig. S5F-G), and observed a strong accumulation of neurons in the intermediate zone with only ∼20.3% neurons entering the cortical plate. Expression of the GPC4^nL^ mutant partially rescued this phenotype, increasing the proportion of neurons entering the cortical plate to ∼40.5%, compared to control conditions (∼75.25%)(Fig. S5H). Taken together, these results show that LRRTM4-GPC4 interactions regulate cortical neuron migration *in vivo* (Fig. 5G).

### LRRTM4-glypican interaction control cancer cell migration via conserved mechanisms

Glypicans are expressed broadly during development, not just the nervous system^55,56^, and are frequently upregulated in tumors^57,58^. Given that LRRTM4 also attenuates GPC3-dependent repulsion in neurons (Figure S6A-B), and that we previously demonstrated a role for the GPC3-Unc5D complex in neuroblastoma cell migration^8^, we investigated whether GPC-LRRTM4 interaction also directs tumor cells. We analyzed the expression of LRRTMs and glypicans in published genome-wide RNA expression profiles of human protein-coding genes in 997 human cancer cell lines^59^ and 32 tumor types from the Cancer Genome Atlas^60^. Among LRRTM family members (LRRTM1-4), LRRTM4 showed the highest expression in both cancer cell lines and related tumor samples (Figure 6A-B). Most glypican family members showed high expression in these datasets, with the exception of GPC5 (Figure S6C-D). As LRRTM4 is predominantly expressed in neural tissues^61^, we measured its prognostic value in tumors such as glioma and neuroblastoma. We found that high LRRTM4 expression correlates with improved patient survival (Figure 6C and S6E-F), whereas GPC4 show the opposite trend, with higher expression associated with poorer survival (Figure 6D).

**Figure 6.**
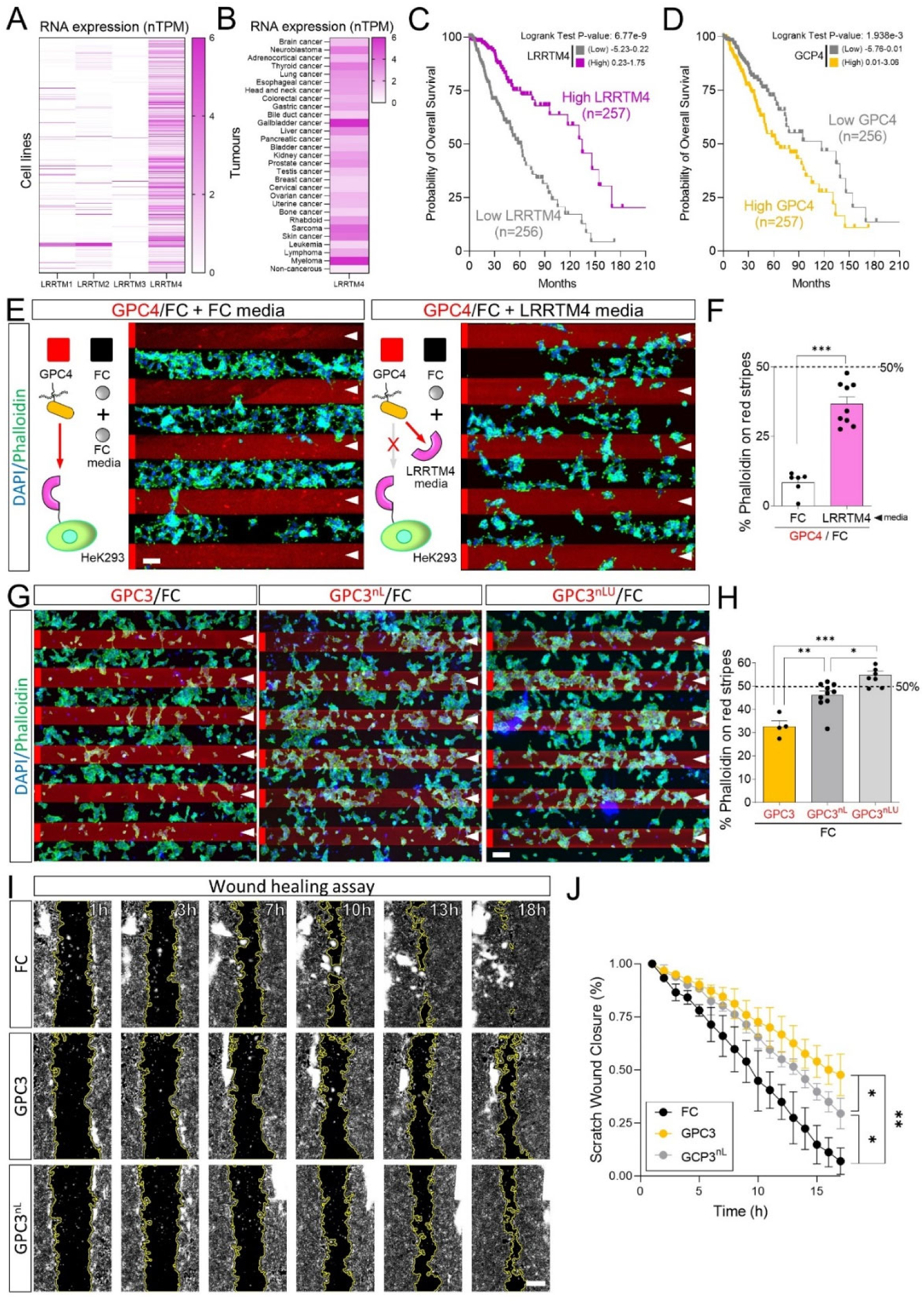
LRRTM4-glypican interaction directs cancer cell migration. (A) LRRTM (1-4) RNA expression in 997 human cancer cell lines, shown as normalized transcripts per million (nTPM). (B) As in panel A, but for 32 tumor types from the Cancer Genome Atlas. (C) Kaplan-Meier survival analysis of LRRTM4 expression in glioma. Patients were stratified into high (n = 257) and low (n = 256) LRRTM4 expression groups. (D) Kaplan-Meier survival analysis of GPC4 expression in glioblastoma, stratified as in C. (E) Dissociated HEK293 cells grown on alternate stripes containing FC (black) and GPC4 proteins (red) in the presence of soluble FC or LRRTM4 ectodomain. Cells were stained with phalloidin (green) and DAPI (nuclei, blue). Red stripes are indicated by white arrowheads. (F) Quantification of the percentage of phalloidin pixels on red stripes. n > 3 independent experiments. ∗∗∗p < 0.001, Student’s t test. (I) Similar to E, but HeK2923 cells were cultured on alternate stripes containing FC (black) and GPC3, GPC3nL or GPC3nLU (red). (H) Quantification of the percentage of phalloidin pixels on red stripes. n > 3 independent experiments. ∗p < 0.05, ∗∗p < 0.01, ∗∗∗p < 0.001, one-way ANOVA test with Tukey’s post hoc analysis. (I) Snapshots from time-lapse imaging of a wound healing assay using HEK293 cells plated on FC, GPC3 or GPC3nL-coated surfaces. The wound area is outlined in yellow. (J) Quantification of the wound closure over time. n=2-3 independent experiments. ∗p < 0.05, ∗∗p < 0.01, one-way ANOVA test with Tukey’s post hoc analysis. Scale bar 100um (E, G), 500um (I).

We next selected several tumor cells lines expressing LRRTM4 (Figure S6G) to assess the potential role of its interaction with glypicans in cell migration. Using stripe assays, we found that GPC4 ectodomain is strongly repulsive for HEK293 and HeLa cells, which was significantly attenuated upon addition of the soluble LRRTM4 ectodomain to the medium (Figure 6E-F and S6H-I).

Moreover, the repulsion of HEK293 cells from GPC3 core domain was reduced when these cells were challenged with the non-LRRTM binding mutant, GPC3^nL^, and was further diminished with the double mutant lacking both LRRTM and Unc5 binding capabilities (GPC3^nLU^), indicating the contribution of both interactions (Figure 6G-H). Finally, we performed a wound healing assay using HEK293 cells plated on control (FC), GPC3, or GPC3^nL^-coated surfaces. In these assays, HEK cells migrated less to close the gap on GPC3-coated surfaces compared with FC controls. This inhibitory effect was partially rescued when using the mutant GPC3^nL.^, confirming that the migration delay is at least partially due to interaction with LRRTM receptors (Figure 6I-J and Movie S3). Taken together, the results support a regulating role of LRRTM-glypican interactions in cancer cell migration.

## Discussion

Cell migration is controlled by complex signaling mechanisms that rely on dynamic interactions between cell-surface receptors and their extracellular ligands. A recurring feature of this process is that the same molecules can elicit distinct cellular responses depending on their binding partners and cellular context. Here, we identify LRRTM-glypican interactions as a new example of such context-dependent signaling, where they promote adhesion during synapse formation, while inducing contact-dependent repulsion during neuronal migration in the developing brain. Moreover, we show that this mechanism extends to tumor cells, where both LRRTM4 and glypicans are frequently upregulated. Together, our findings support an emerging view that conserved cellular mechanisms operate across distinct biological contexts, including brain development and tumor dissemination.

In the developing brain, we found that LRRTMs are among the most highly expressed LRR-containing receptor families, consistent with a potential role in early developmental processes, which precede synapse formation. Indeed, we find that LRRTMs direct the migration of cortical neurons, possibly by regulating their interaction with the radial glia scaffold. Mechanistically, LRRTM4 is enriched in migrating neurons and binds glypicans *in trans* on opposing RG cells, whose processes guide neurons to the cortical plate. This configuration mirrors that proposed for synapse formation: LRRTM4 on the postsynaptic site interacts *in trans* with glypicans on the presynaptic side^19,20^. This is reminiscent of other LRR protein families, such as FLRTs, in which the same the same receptor interactions operate in both synapse formation and neuronal migration^12,16^. Interestingly, the same trans-interaction that promotes adhesion at synapses, can instead mediate repulsion during cell migration. Such dual roles in adhesive synaptogenesis and repulsive cell guidance have been described for FLRT and Eph/ephrin signaling^12,62,63^, as well as Teneurins^64^, although the mechanisms underlying this switch remain unclear. Other LRRTM configurations are conceivable, as observed for FLRT proteins, which can engage in both homo- and heterophilic interactions between neurons^11,23,65^. For example, glypicans such as GPC1 and GPC6, as well as neurexins, are expressed in migrating neurons, suggesting that transient neuron-neuron interactions may occur during migration. Our results are a starting point for exploring alternative interaction topologies and determining whether cis-interactions modulate these responses^64^.

Our findings extend LRRTM-glypican interactions beyond the nervous system to tumour cell migration. Glypicans are frequently upregulated in tumors^57,58^, and their high expression correlates with poor prognosis^30,66^, as also indicated by our analysis for GPC4 in glioma and neuroblastoma (Fig. 6A-D). In contrast, among LRRTM family members, LRRTM4 is also expressed in several tumors but shows an opposite trend, with higher expression correlating with improved patient survival. These observations raise the possibility that LRRTM4 may act as a tumor suppressor in glypican-expressing tumors. This interpretation is consistent with the repulsive signaling elicited by LRRTM–glypican interactions, which reduces tumor cell migration and may thereby limit tumor progression and dissemination. A similar scenario has been described for EphB receptors, which regulate intestinal epithelial architecture through repulsive interactions with ephrin-B ligands^67^. Loss of this repulsion, due to reduced EphB expression, accelerates tumor invasion and progression^67^.

We previously showed that GPC3 forms an octameric complex with Unc5D via glycan–glycan interactions involving N-linked glycans on GPC3 and C-mannosylated tryptophans on Unc5D^8^. This complex regulates both neuron migration, where GPC3 on RG cells induces repulsion of Unc5D-expressing neurons, and neuroblastoma dissemination. Interestingly, in stripe assays neurons were repelled by both GPC3 and a mutant unable to bind Unc5D^8^, indicating that additional GPC3 receptors mediate repulsion independently of Unc5D. Our results point to LRRTMs as candidate receptors mediating this effect, at least in HEK293 cells, as a GPC3 mutant defective in both Unc5D and LRRTM binding capabilities has a nearly neutral effect in stripe assays. Our AlphaFold-based structural predictions, validated with mutant proteins and cell-based assays, show that LRRTMs bind directly to the glypican core domain through their LRR region. This contrasts with previous studies indicating that glypican binding to LRRTM4, but not LRRTM2, requires HS chains. These conclusions were based on competition experiments in which excess HS reduced LRRTM4 binding to GPC4^20^, as well as enzymatic removal of HS or the use of glypican mutants lacking glycosaminoglycan attachment sites^61^. One possible explanation for this apparent discrepancy is that HS, whilst not essential for the interaction, still contributes to LRRTM binding to glypicans. Consistent with this idea, HS has been shown to influence LRRTM interactions with other binding partners. For example, recent studies indicate that HS modulates the interaction between LRRTM and neurexins, as mice expressing a mutant form of LRRTM4 that cannot bind HS show reduced levels of neurexin in the dentate gyrus^68^. Moreover, HS has been shown to enhance neurexin-1 function, as mice lacking HS on neurexin-1 show reduced survival and impaired synapse formation^69^. Together, these observations suggest that HS may expand the binding repertoire or stabilize interactions of glypicans, thereby fine-tuning receptor–glypican binding. Similarly, glypicans are known to bind to Wnt and Frizzled receptors using a combination of HS-mediated and core domain interactions, which can accommodate also lipid binding^40,70^. In future work, it would be interesting to assess the possible role of HS chains functioning in LRRTM-GPC mediated regulation of cell migration and to understand how these receptors impact on other GPC- and LRRTM-mediated interactions. Moreover, because Wnt signaling is active in radial glial cells during cortical development, where it regulates both neurogenesis and neuronal migration^71,72^, it will be also interesting to explore whether LRRTMs function within broader glypican signaling assemblies, including Wnts and Unc5 receptors.

In conclusion, our study identifies LRRTM–glypican interactions as a mechanism regulating cell migration across distinct biological contexts. In the developing cortex, glypicans presented by radial glial cells act through LRRTM4 to control neuronal migration by repulsive signaling, while the same interaction similarly influences tumor cell migration. These findings illustrate how protein complexes previously linked to synapse development also participate for cell navigation and reveal a conserved mechanism shared between developmental and tumor migration. Given the widespread expression of LRRTMs and glypicans, this mechanism may also operate in additional biological contexts.

## Acknowledgements

We thank Ana López (María de Maeztu Unit of Excellence, Institute of Neurosciences, UB, CEX2021-001158-M) for animal colony management. Maria Calvo from the Advanced Microscopy service (CCiT, UB) for help with confocal microscopy. We thank the proteomics facility of the Max Planck Institute of Biochemistry (RRID:SCR_025745; Martinsried, Germany) and its facility members for support and sample processing. C.P was funded by an FI fellowship from Generalitat de Cataluña and the Wellcome Trust (226647/Z/22/Z).M.B.-S was funded by a Pelly-Bannister Scholarship (Somerville College, Oxford). V.C. was supported by the ERC Synergy grant “SUNRISE” (101167188). E.S. was supported by a Wellcome Trust Discovery Award “MIGRATE” (226647/Z/22/Z) and the ERC Synergy grant “SUNRISE” (101167188). D.d.T was funded by the Wellcome Trust (226647/Z/22/Z) and the ERC Synergy grant “SUNRISE” (101167188). Part of the glioma cohort analysis results were generated by the TCGA Research Network: https://www.cancer.gov/tcga

## Author contributions

C.P. performed initial binding studies, most of the in vitro experiments, led the in vivo work and performed expression analyses and co-conceptualized the work. M.B.-S. performed structural predictions, protein engineering, protein production, cell-binding and cell aggregation assays and co-conceptualized the work. M.C.-O. produced plasmids and protein. I.R.-N. performed western blots. K.e.O. assisted with the structural analysis. H.Y.L. contributed to glioma analysis, G.S.-B contributed to pull down experiments, R.K. oversaw the developmental studies. V.C. contributed to neuroblastoma analysis. E.S. oversaw structural biology, binding studies and protein engineering and co-conceptualized the work. D.d.T. oversaw the in vivo work, expression analyses and stripe assays and oversaw the conceptualization of the work. All authors have contributed to the manuscript.

## Methods

### Data and code availability

- Mass spectrometry data has been deposited in the PRIDE repository, where it will be publicly available as of the date of publication.
- This paper does not report original code.
- Any additional information required to reanalyze the data reported in this paper is available from the lead contacts Elena Seiradake, and Daniel del Toro

### Material availability

This study did not generate new unique reagents.

## Experimental Model and Subject Details

### Mouse embryos

All mice (C57BL/6 background) were housed with a 12h:12h light:dark cycle and food/water available ad libitum. All animal experiments were used in accordance with the ethical guidelines (Declaration of Helsinki and NIH, publication no. 85-23, revised 1985, European Community Guidelines, and approved by the local ethical committee (University of Barcelona, 225/17 and Generalitat de Catalunya, 404/18).

### Primary cultures

Neurons were dissociated from cortices of E15.5 embryos and cultured on stripes. Neurons were cultured for 1 days in vitro at 37°C, 5% CO2 in Neurobasal medium supplemented with B27. Neurons were used for stripe assays and were fixed with 4% Paraformaldehyde for 10 min followed by immunostaining.

### Cell lines

K562 suspension cells (ATCC; CCL-243; RRID: CVCL_0004) were cultured in RPMI-1640 media (LGC Standards, Cat#ATCC 30-2001) supplemented with 10% FBS (GIBCO, Cat#10437028). K562 cells were maintained in sterile conditions in a 37°C, 5% CO2-incubator. Expi293F cells (ThermoFischer Scientific, Cat#A14527; RRID: CVCL_D615) were cultured in Expi293 Expression medium and maintained in sterile conditions in a 37°C, 70% humidity, and 120 rpm 8% CO2-incubator. HEK293 and HeLa cells were cultured in DMEM (Life Technologies) supplemented with 10% FBS and 5% L-Glutamine. Cell lines were maintained in sterile conditions in a 37°C, 5% CO2-incubator.

## Method Details

### Lectin pull-down

Pull-down experiments were performed as previously described^12^. Fresh E15.5 mouse cortices were homogenized for 1 min at 4°C with an electric homogenizer using the following lysis buffer: 50 mM Tris-HCl (pH 7.4), 150mM NaCl, 2mM EDTA, 1% Triton X-100 and protease inhibitors (Sigma-Aldrich 04693132001). Samples were incubated on ice for 30 min and centrifuged for 10 min at 3000 rpm. Supernatant was collected and protein was measured using the Bio-Rad protein assay (Biorad, 5000001). 1 mg of protein at a final concentration of 2 mg/ml in lysis buffer (volume: 500 ml) was used for each pull-down. Control pull-down contained lysate and 1 mg of goat anti-human IgG antibody (Jackson Immunoresearch,109-005-098) while Lectin pull-down used 1 mg of Lectin from Triticum vulgaris (L1882). Samples were incubated overnight at 4°C under rotatory agitation. Then samples were centrifuged for 5 min at 3000 rpm and washed three times (first wash with 400 ml of lysis buffer, second wash with 1:1 (v/v) lysis buffer:PBS, last wash only PBS). Pulled-down samples were processed for mass spectrometry (MaxQuant run, Proteomic facility, Max Planck Institute of Biochemistry, Martinsried, Germany).

### Analysis of published single cell RNASeq dataset

Single RNAseq data for cortex samples were obtained from the published NCBI Gene Expression Omnibus with accession number GSE15316443. We used the same UMAP coordinates and metadata information with the cluster categorization provided by the authors^73^. We used the R package Seurat (v.4.0.1) to perform all the analysis.

Glioma cohort analysis data were obtained from The Cancer Genome Atlas low-grade glioma (TCGA-LGG) dataset and analyzed using cBioPortal^74–76^. Neuroblastoma cohort analysis data were obtained from the SEQC-498 cohort (GSE49710) and analyzed using the R2 Genomics Analysis and Visualization Platform (https://hgserver1.amc.nl). Expression levels of LRRTM1-4 and GPC4 were stratified into high- and low-expression groups according to the median expression value. The correlation between gene expression and overall survival was assessed by Kaplan-Meier survival analysis. The resulting data were further exported and plotted using GraphPad Prism 10.

### RNA In situ hybridization (ISH)

Embryonic brains were fixed in 4% PFA over-night. 10 μm Cryo-sections were pre-treated using the RNAscope Universal Pretreatment Kit (Advanced Cell Diagnostics, Cat#322380). RNA In Situ Hybridizations (ISH) were performed using the RNAscope Fluorescent Multiplex Reagent Kit (Advanced Cell Diagnostics, Cat#320850) according to manufacturer’s instructions. Images were acquired using a Zeiss LSM880 confocal laser scanning microscope and processed with ImageJ software.

### Binding assay

Hela cells were passed to a 6 well plate containing covers. Next day, cells were transfected with Fugene (Promega, E2311) to overexpress our protein of interest. In this case, the construct used were: PCAGCIG-GPC4 WT and mutant. Next day, His-tagged LRRTM4 protein from R&D (#5377-LR) were pre-clustered with 6x His Tag Alexa Fluor647-conjugated Antibody (#IC050R) for 30 min at RT in HBSS. Then a drop of this mix was added to each cover for 20 min on ice. Cells were washed with phosphate buffer saline (PBS) and fixed in 4% PFA (20 min, on ice). Then cells were washed with PBS and mounted with DAKO (SoleyBio, S1964). DAPI was used for nuclei staining. Images were acquired using a Zeiss LSM880 confocal laser scanning microscope and processed with ImageJ software.

### Time-lapse experiments

To perform the time-lapse analysis of cortical neurons migrating on surfaces coated with FC (control), GPC4 or LRRTM4, cortical explants from E15.5 mouse embryos were cultured on 60-mm dishes that were coated for 30 min with 50 μg/ml of each protein in PBS. After 30 min, the dish was washed with PBS and coated with 20 μg/ml Laminin in PBS for at least 2 hours. The dish was next washed with PBS and cortical explants (E14.5) were cultured. After 2 hours in culture, samples were imaged with an Optical microscope: the Widefield Leica AF6000 that was equipped with a temperature-controlled carbon dioxide incubation chamber set to 37°C, 65% humidity and 5% CO2. Sequential images were acquired every 10 min and for 8 h. Analysis and tracking of all the neurons exiting the explant was carried out in ImageJ using the semi-automatic Trackmate pluggin.

### Nanofiber experiments

6 well-plate parallel nanofibers (700nm width, Sigma, Z759333-1EA) were coated with 40 μg/ml of specified protein (FC, GPC4 or LRRTM4) and 100 μg/ml poly-D-lysine (Sigma) in PBS overnight (37°C, 65% humidity and 5% CO2). Next day, plates were washed with PBS and coated with 20 μg/ml laminin in PBS overnight. The next day, plates were washed with PBS and 36 cortical explants (E14.5) were placed per well and cultured for 2 days in Neurobasal medium supplemented with B27 (Invitrogen) supplemented with 0.4% methyl-cellulose (Sigma). Then the explants were fixed with 4% PFA for 20min and washed with PBS. Nuclei were counterstained with DAPI before mounting. Mosaic images of each well were taken with a Widefield Leica AF6000 microscope. Analysis of the distance covered by cells labeled with DAPI was done in a semi-automatic mode using a custom ImageJ macro.

### Stipe assays and immunostaining

We prepared the stripe assays essentially as previously described^12^. 50 μg/ml of Fc recombinant protein, GPC4, LRRTM1-4, GPC3, GPC3nL or GPC3nLU were mixed with Alexa647-conjugated anti-hFc antibody (Thermo Fisher Scientific) in PBS. Proteins were injected into matrices (90 μm width) (17546017) and placed on 60 mm dishes, resulting in red fluorescent stripes. After 30 min incubation at 37°C, dishes were washed with PBS and matrices removed. Dishes were coated with 50 μg/ml Fc protein mixed with 150 μg/ml anti-hFc (Jackson ImmunoResearch, cat#62-8400) for 30 min at 37°C and washed with PBS. Stripes were further coated with 20 μg/ml Laminin in PBS overnight and washed with PBS next morning. Cortical neurons (E15.5) were cultured on the stripes in Neurobasal medium supplemented with B27 (Gibco). After 24 hours neurons were fixed with 4% PFA in PBS for 10 min at room temperature (RT). Neurons were washed and incubated with rabbit monoclonal anti-beta-III tubulin antibody (Sigma-Aldrich) after 10 min permeabilization in 1% BSA, 0.1% Triton X-100 in PBS. Alexa488 anti-rabbit IgG secondary antibody (Thermofisher, cat#A-21206) was used to visualize the tubulin signal. Nuclei were counterstained with DAPI before mounting. The numbers of beta-III-tubulin-positive (green) pixels on red or black stripes were quantified with ImageJ (version 2.9.0/1.53t)63 using a custom-made automatic macro that is available upon request. Additionally, on top of some stripes, we cultured cortical explants (E15.5) for 48h to study the axons behavior. For explants culture the neurobasal medium was supplemented with B27 (Invitrogen) and 0.4% methyl-cellulose (Sigma). For cell lines experiments, laminin coating is not needed. After stripes making 30000cells/stripes (in 150uL volume) were seeded in DMEM with 2% FBS. After 6 hours cells were fixed with 4% PFA in PBS for 10 min at room temperature (RT). Cells were washed and incubated with green conjugated phalloidin (A12379) 1:400 for 30 min after 10 min permeabilization in 1% BSA, 0.1% Triton X-100 in PBS. Nuclei were counterstained with DAPI before mounting.

### Cell surface expression tests

HEK293 cells were transiently transfected with transmembrane constructs of GPC4 and LRRTM4 (WT and mutants) as explained above in the binding assay. All constructs were fused to an extracellular HA-tag (for LRRTM4) or Flag-tag (for GPC4). Depending on the tag included, anti-HA (SIGMA-Aldrich, Cat No. H3663) or anti-Flag (SIGMA-Aldrich, Cat No. F1804) were previously incubated with an Alexa Fluor 647 antibody (thermofisher) (1:4) in HBSS and and added to the cells at a concentration of 5 μg/ml. Cells were incubated for 30 min on ice and washed with PBS. Cells were fixed in 4% PFA (20min, on ice) and mounted. DAPI was used as nuclei staining and Images were acquired using a Zeiss LSM880 confocal laser scanning microscope. Non-transfected cells acted as negative control.

In case of experiments for sh validation, HEK293 cells were transiently co-transfected with transmembrane constructs of the WT version of GPC4 or LRRTM4 and the corresponding shRNA (see IUE for sequence information) embedded in the in the pCAG-miR30 vector (1:3).Then, after 24h of incubation, some of the cells were used to test the surface expression of the proteins of interested by the corresponding tag as described above and the rest of the cells were lisated using the following lysis buffer: 50 mM Tris-HCL (pH 7.4), 150mM NaCl,2mMEDTA,1%Triton X-100 and protease inhibitors (Roche ref. 04693116001). Samples were then incubated on ice for 20 min and centrifuged for 10 min at 3000 rpm. Supernatant was collected and protein was measured using the Bio-Rad protein assay (Biorad, 5000001) and Western Blot was performed to quantified the knockdown of the protein of interested.

### Western Blot

Supernatant proteins (15 ug) from the cells lysates were loaded in SDS–PAGE and transferred to nitrocellulose membranes (GE Healthcare, LC, UK). Membranes were blocked in TBS-T (150 mM NaCl, 20 mM Tris-HCl, pH 7.5, 0.5 ml Tween 20) with 5% BSA and 5% non-fat dry milk. Immunoblots were incubated overnight at 4° with anti-FLAG at and anti-HA at 1:1000 in PBS with 0.2% Sodium Azide. After three washes in TBS-T, blots were incubated for 1 h at room temperature with anti-rabbit horseradish peroxidase-conjugated secondary antibody (1:2000; Promega, Madison, WI, USA) and washed again with TBS-T. Immunoreactive bands were visualized using the Western Blotting Luminol Reagent (Santa Cruz Biotechnology). For loading control, a mouse monoclonal antibody for actin was used (1:100.000, Sigma Aldrich, #A3854).

### In utero electroporation

In utero electroporation was performed at E13.5/E15.5 with anesthetized C57BL/6 mice as previously described^12^. DNA plasmids were used at 2 μg/μl and mixed with 1% fast green (Sigma-Aldrich, final concentration 0.2%). Plasmids were injected into the ventricle with a pump-controlled micropipette. After injection, six 50 ms electric pulses were generated with electrodes confronting the uterus above the ventricle. The abdominal wall and skin were sewed, and the mice were kept until E16.5/E18.5 embryonic stage. To knockdown GPC4 and LRRTM4 by shRNA in vivo, we used two different shRNAs embedded in the in the pCAG-miR30 vector, with the following sequences: #shRNAGPC4 (sequence: GCGAACAGTGCAACCATTTGC) and #shRNALRRTM4 (sequence: GTCACAAGGTTTATCCTTAA).

Additionally, DNA plasmids to overexpressed WT or mutant forms of the proteins were designed using SnapGene Viewer and validated in Heks293 cells.

### Morphological analysis

ShRNA plasmids to KO GPC4 and LRRTM4 were electroporated at E13.5. After 3 days, embryonic brains were collected, fixed in 4% PFA overnight and vibratome cut into 75mm sections. Single cell morphology was reconstructed and analyzed using ImageJ (version 1.53) as described previously^23,77^. For single cell morphology analysis in the lateral cortex was used after maximum projection of a z stack representing 50-60um (one image per 5um). Single cell morphology from GFP-expressing neurons was reconstructed and analyzed using ImageJ (version 1.49). 15–20 neurons per cortical layer (upper CP, lower CP and IZ) were quantified per section (2-3 sections per brain and three independent brains per condition).

### Wound Healing Assay

Wound healing assays were performed on surfaces coated with FC (control), GPC3, or GPC3nL. Twelve-well plates were coated with 50 µg/mL of each protein in PBS for 30 min at RT. After coating, µ-Slide 2 Well Culture-Inserts (Ibidi, Cat. No. 80209) were carefully placed in each well. Then Hek293 cells were suspended at a density of 0.5 × 10⁶ cells/mL, and 70 µL of the cell suspension was added to each side of the insert. Cells were cultured in DMEM supplemented with 2% FBS and incubated at 37 °C with 5% CO₂. Cells were allowed to reach confluency (∼24 h). After monolayer formation, the silicone inserts were gently removed, and additional medium was added. Wound closure was monitored using the Opera Phenix high-content imaging system over 18 h, with images captured every hour. Wound closure was quantified using the wound healing size tool from ImageJ^78^, calculating the decreasing percentage in wound area over time.

### Vectors and cloning

We cloned constructs of mouse LRRTM1 (Uniprot: Q8K377) (full length: residues 37-522), mouse LRRTM2 (Uniprot: Q8BGA3) (full length: residues 34-515; ecto: 34-421; nG mutant: H211N+E213T), mouse LRRTM3 (Uniprot: Q8BZ81) (full length: residues 31-582), mouse LRRTM4 (Uniprot: Q80XG9) (full length: residues 31-590; ecto: 31-424; nG mutant: H211N+E213T), human GPC3 (nL mutant: V109N+Q111T; nLU mutant: V109N+Q111T+N241Q), human GPC4 (full length, residues 24-556; nL mutant: K104N+D106T), human GPC5 (full length, residues 25-572), mouse Nrx1ß (Uniprot: P58400) (full length: residues 50-472). We have used previously published hGPC3 (core), rUnc5B (full length), rUnc5DIg1-Ig2-Tsp1, mLphn1 (Lec), and mLphn2 (Lec) constructs/boundaries as indicated^8,12^.

For protein expression, the ectodomain constructs were cloned into the Age1-Kpn1 site of the pHL-Sec vector (Addgene; Cat#99845). This pHL-Sec vector that contains an N-terminal secretion signal peptide which is used instead of the native signal peptide of the proteins. The vector also contains a 6xHis tag and an eAvi-tag (protein sequence: GLNDIEAQKIEWHE) at the C-terminus. For cell binding, cell-based aggregation assays and functional assays, full length constructs were used, cloned into the Xho1-Not1 site of the pCAGIG vector (Addgene; Cat#11159). pCAGIG was modified to express Tomato instead of GFP, for certain experiments, and is then referred to as pCAGIT. LRRTM, Nrx1ß, and Unc5B constructs were cloned to contain an HA tag at the N-terminus (protein sequence: YPYDVPDYA) in the pCAGIT or pCAGIG vector, whilst the GPC constructs contained a FLAG tag at the N-terminus (protein sequence: DYKDDDDK) in the pCAGIG vector.

### Alphafold structural predictions

For the structural prediction of the LRRTM4-GPC4 complex we used Alphafold2 multimer with default settings. The following sequences were inputted into Alphafold2 as a FASTA file:

Mouse LRRTM4 (Q31-H423): Uniprot ID-Q80XG9

RACPKNCRCDGKIVYCESHAFADIPENISGGSQGLSLRFNSIQKLKSNQFAGLNQLIWLYLDHNYISSVDEDAFQGIRRLKELILSSNKITYLHNKTFHPVPNLRNLDLSYNKLQTLQSEQFKGLRKLIILHLRSNSLKTVPIRVFQDCRNLDFLDLGYNRLRSLSRNAFAGLLKLKELHLEHNQFSKINFAHFPRLFNLRSIYLQWNRIRSVSQGLTWTWSSLHTLDLSGNDIQAIEPGTFKCLPNLQKLNLDSNKLTNVSQETVNAWISLISITLSGNMWECSRSICPLFYWLKNFKGNKESTMICAGPKHIQGEKVSDAVETYNICSDVQVVNTERSHLAPQTPQKPPFIPKPTIFKPDAVPATLEAVSPSPGFQIPGTDHEYEHVSFH

Mouse GPC4 (K26-E536): Uniprot ID-P51655

KSKSCSEVRRLYVSKGFNKNDAPLYEINGDHLKICPQDYTCCSQEMEEKYSLQSKDDFKTVVSEQCNHLQAIFASRYKKFDEFFKELLENAEKSLNDMFVKTYGHLYMQNSELFKDLFVELKRYYVAGNVNLEEMLNDFWARLLERMFRLVNSQYHFTDEYLECVSKYTEQLKPFGDVPRKLKLQVTRAFVAARTFAQGLAVARDVVSKVSVVNPTAQCTHALLKMIYCSHCRGLVTVKPCYNYCSNIMRGCLANQGDLDFEWNNFIDAMLMVAERLEGPFNIESVMDPIDVKISDAIMNMQDNSVQVSQKVFQGCGPPKPLPAGRISRSISESAFSARFRPYHPEQRPTTAAGTSLDRLVTDVKEKLKQAKKFWSSLPSTVCNDERMAAGNENEDDCWNGKGKSRYLFAVTGNGLANQGNNPEVQVDTSKPDILILRQIMALRVMTSKMKNAYNGNDVDFFDISDESSGEGSGSGCEYQQCPSEFEYNATDHSGKSANEKADSAGGAHAE

The resulting best ranked prediction was visualised using ChimeraX and Pymol. For the GPC1-6 monomer predictions we inputted in Alphafold3 the following sequences from Uniprot: GPC1 (P35052), GPC2 (Q8N158), GPC3 (P51654), GPC4 (O75487), GPC5 (P78333), and GPC6 (Q9Y625). For the LRRTM1-4 monomer predictions we inputted in Alphafold3 the following sequences from Uniprot: LRRTM1 (Q8K377), LRRTM2 (Q8BGA3), LRRTM3 (Q8BZ81), and LRRTM4 (Q80XG9).

### ConSurf and multiple sequence alignments

To produce the conservation scores for the structures we used ConSurf Colab (https://colab.research.google.com/drive/1PhDXX7k12oUsV6T_xkXC3Rm9R99e7tHz#scroll To=zBTwgYcIK6Ul). For the ConSurf run we used the standard settings, but we inputted a custom MSA using Clustal Omega. For this, we used the following sequences for LRRTM4 from Uniprot: A0AB13AAX7 (Danio rerio), A0A6I8PRN2 (Xenopus tropicalis), A0A5F8H6H0 (Monodelphis domestica), F1NP47 (Gallus gallus), Q86VH4 (Homo sapiens), and Q80XG9 (Mus musculus); and the following sequences for GPC4 from Uniprot: B3DJ94 (Danio rerio), F7CR54 (Xenopus tropicalis), A0A5F8GD71 (Monodelphis domestica), A0A8V0Z7Y7 (Gallus gallus), O75487 (Homo sapiens), and P51655 (Mus musculus).

The conservation scores were mapped onto the structures using Pymol and the ConSurf provided script.

### Protein expression and purification

For the purification of all proteins used in this study (except the ones described at the end of this section) we transfected 120 ml of Expi293F at a density of 3×106 cells/ml with 120 μg of DNA and 360 μl of PEI MAX (Polysciences, Cat#24765-1) in a 1:3 DNA:PEIMAX ratio following the manufacturer protocol. Cells were left to express the protein for three days and then the media were harvested, clarified by centrifugation and filtered using 0.22 μm sterile filters (Starlab; Cat# S1120-8810). The filtered media was concentrated and buffer-exchanged to diafiltration buffer (1X PBS (Invitrogen; Cat#3002), 20 mM Tris-HCl pH=7.5, 150 mM NaCl). Then this media was passed through a 5 ml HisTrapTM HP column (Cytiva, #Cat17-5248-02) at a flow of 5 ml/min. The column was washed with wash buffer (20 mM Tris-HCl, 300 mM NaCl, 40 mM Imidazole (Sigma-Aldrich, Cat#I3386)) and the protein eluted with 20 mM Tris-HCl, 300 mM NaCl, 500 mM Imidazole. 450 μl of the centre peak fraction was then injected into a pre-equilibrated SuperdexTM 200 Increase 10/300 GL (Cytiva; Cat#28-9909-44) in 20 mM Tris-HCl (SIGMA-Aldrich; Cat#7365-45-9) (pH=7.5) and 200 mM NaCl running buffer. Elution fractions were collected, analysed using SDS-PAGE and the peak fractions were frozen at −80°C until use. rUnc5DIg1-Ig2-Tsp1, mLphn1Lec, mLphn2Lec were used from previous publications^8,12^

### Pull-down of LRRTM-GPC

Pull-down experiments to investigate the LRRTM – GPC interaction were performed by coupling 0.2 μM of biotinylated hGPC3core to high-capacity streptavidin agarose resin (Thermo Fisher Scientific, Cat#20357). After incubation of the beads with the biotinylated protein for 1 h at RT, 0.4 μM of the soluble protein (mLRRTM2ecto, rUnc5DIg1-Ig2-Tsp1, mLphn2Lec, or hLRRTM4 (RnD systems, Cat#5377-LR) in 20mM Tris-HCL, pH 7.5, 200mM NaCl, 1% BSA was incubated with hGPC3core-coated strep beads for 2 h. Beads were washed with 20mM Tris-HCL, pH7.5, 200mM NaCl and the proteins were eluted with SDS-containing gel-loading buffer supplemented with 5% beta-mercaptoethanol. Samples were loaded into a 4-12% NuPAGE Novex Bis-Tris gels (Thermo Fischer Scientific; Cat#ENP0336BOX), run for 45 mins at 180 V. The gels were transferred in the NuPAGE® XCellIITM Blot transfer module for 90 minutes at 30 V onto a nitrocellulose membrane (Cytiva, Cat#10600013), sandwiched with two blotting papers and five sponges, submerged in 1X NuPAGE® Transfer Buffer (InvitrogenTM, Cat#NP00061) supplemented with 20% ethanol. After transferring, the membrane was blocked for 30 minutes using 3% BSA (w/v) (InvitrogenTM, Cat#A7906) in PBST (1X PBS with 0.1.% Tween20) and washed with PBST only for 30 mins. Then the membranes were incubated in the primary antibody (mouse anti-His (Qiagen, Cat#34660) prepared in PBST with 3% BSA in a 1:1000 dilution) solution for 1 hour. After incubation, the membrane was washed for 30 minutes with PBST and incubated for 1 hour with the secondary antibody solution (anti-Mouse IgG HRP (Sigma-Aldrich, Cat#A0168) diluted 1:10000 in PBST with 3% BSA). Then the membrane was washed with PBST for 45 minutes and developed with the ECLTM Western blotting detection reagents (Cytiva, Cat#RPN2106) and visualised using AmershamTM HyperfilmTM ECLTM films (Cytiva, Cat#28-9069-35).

### Cell-cell aggregation assay

K562 suspension cells were cultured in RPMI-1640 media (no phenol red) (Invitrogen; Cat#11835030) supplemented with 10% FBS and 5% L-Glutamine. The cells were harvested by a 3 min spin at 200g, washed with PBS, spined again and resuspended in R buffer (Neon Transfection System 100 μL Kit; Thermo Fischer Scientific; Cat#MPK10025). Cells at a concentration of 2×107 cells/ml were transfected with 15 μg of control pCAGIG/pCAGIC plasmids, or those coding for LRRTM1-4, Nrx1ß, Unc5B or GPC3-5 constructs using the Neon transfection system for electroporation (Settings: 1450V, 3 pulses, 10 ms). Eighteen hours after transfection, cells were harvested and used at a concentration of either 2 x 105 cells/ml or 4 x 105 cells/ml in aggregation media (Neurobasal-A media without phenol red (Thermo Fischer Scientific; Cat#12349015) supplemented with 2 mM L-glutamine (Life Technologies; Cat#25030-024), 10% FBS, 4% B-27 and 20 mM HEPES) in a 24-well plate. Cells were then left to aggregate at 37°C, 5% CO2 and 250 rpm for 90 minutes. After the incubation, the cells were imaged directly in the 24-well plate after a slight shake using a Nikon ECLIPSE TE2000-U inverted fluorescence microscope. The total area of cells and the total area of the aggregates for each picture were calculated using the Analyze particle tool in ImageJ. The threshold used to distinguish cells and aggregates was determined at 1284 μm2 (>3/4 cells). For the statistical analysis we used a one-way ANOVA test, with a Tukey’s post-hoc test in GraphPad Prism (version 10 for MacOS, GraphPad Software, San Diego, California USA, www.graphpad.com). Significance was determined when p < 0.05. Pictures showed in Supplementary Fig. 4 were imaged using a THUNDER Imager (Leica).

For the competition aggregation with either hGPC3core or mLphn1Lec, the experiment was performed as stated above but 25 μg of either protein was added to the cells after plating. These were then left to aggregate at 37°C, 5% CO2 and 250 rpm for 90 minutes. After the incubation, the cells were imaged as explained above.

### Cell surface expression in K562 cells

For surface staining, K562 cells were harvested eighteen hours after electroporation (performed as explained above) and cooled to 4°C for 30 mins. Cells were spined at 200g for 3 mins (4°C) and the media aspirated. This process of spinning and aspirating was performed in between all following steps. All the steps in this protocol are performed on non-permeabilised cells, so that the antibody only detects protein that has been trafficked to the cell surface where the tag is exposed to the antibodies. We also perform all steps at 4°C to abolish endocytosis. The cells were then incubated on ice with blocking buffer: HBSS with 1% BSA and 10 mM HEPES (pH 7.5) for 30 minutes on a shaker. Cells were incubated for 1 hour on ice on a shaker after being added the clustered antibodies. For this, we used anti-HA or anti-FLAG (both mouse: SIGMA-Aldrich, Cat#H3663, AB_262051; SIGMA-Aldrich, Cat# F1804-200UG, AB_262044) antibody that was pre-clustered with secondary anti-Mouse antibody-Cy3 or A488 (Invitrogen; Cat#A10521; AB_10373848; Cat#A-11001; AB_2534069) depending on the vector used for 60 minutes (ratio 1:7.5 for primary:secondary antibody) at room temperature in the dark. Cells were washed with PBS, fixed with 4% PFA for 20 minutes, washed with PBS with ammonium chloride and stained with DAPI (0.1 μg/ml) for five minutes on ice and shaking. Cells were then resuspended on 20 μl Immu-Mount and put on the centre of the slide. Finally, a rectangular coverslip was put on top of the resuspended cells. The cells were imaged on a using a Nikon ECLIPSE TE2000-U inverted fluorescence microscope (20X magnification). Colocalisation for the surface quantification was performed using ImageJ with the colocalisation tool and a macro that can be made available upon request. This colocalisation analysis compares the pixels in both channels (green and red) and determines which pixels have a signal in both channels. A ratio of 10% between red and green channels was used. As the cells are non-permeabilised, the signal coming from the red channel (Cy3) can only come from the HA/FLAG-tagged proteins on the cell surface, if present. The area of the pixels that are colocalised between both channels is the normalised by the total transfected cell area. For the statistical analysis we used a one-way ANOVA test, with a Tukey’s post-hoc test in GraphPad Prism (version 10 for MacOS, GraphPad Software, San Diego, California USA, www.graphpad.com). Significance was determined when p < 0.05. Pictures showed in Supplementary Fig. 4 were imaged using a THUNDER Imager (Leica).

## Quantification and statistical analysis

Statistical analyses were performed using GraphPad Prism, employing a two-tailed unpaired Student’s t test (Figures 2C,F,H,J, 6F,H and S6B,H) or Logrank test analysis (Figures 6C,D) when comparing two groups or multiple groups distribution, and one-way ANOVA test with Tukey’s post hoc analysis when comparing multiple groups (Figures 4A,B,C, 5D,F, 6J, S2C,E, S4B,D and S5H) p values represent *p<0.05, **p<0.01, ***p<0.001 and ****p<0.0001. All data are presented as the mean ± SEM, whisker plots or dot plots. All sample sizes and definitions are provided in the figure legends.

## Supplemental item legends

**Supplemental movie S1:** Time-lapse analysis of cortical neurons migrating from E15.5 mouse cortical explants cultured on dishes coated with FC (control) or GPC4 (ectodomain). Explants were imaged 2 h after plating and images were acquired every 10 min for 8 h. Neurons exiting the explant were tracked using the semi-automatic TrackMate plugin in ImageJ.

**Supplemental movie S2:** Time-lapse analysis of electroporated cortical neurons migrating on nanofibers. We electroporated mouse embryos at E13.5 with pCAGIG and peformed explant cultures from the cortex 2 days later (E15.5). After 4 hours in culture, the explants were imaged with a Zeiss Axiovert 200M microscope equipped with a temperature-controlled carbon dioxide incubation chamber set to 37°C, 65% humidity and 5% CO2. Illumination was provided by an X-Cite lamp (series 120, Lumen Dynamics Group), and images were recorded by a Coolsnap HQ camera (Photometrics). Sequential images were acquired every 6 min.

**Supplemental movie S3:** Representative time-lapse imaging of HEK293 cells in a wound healing assay on surfaces coated with FC (control), GPC3, or GPC3nL. Images were acquired using the Opera Phenix high-content imaging system over 18 h, with images captured every hour. Wound closure was quantified using the wound healing size tool from ImageJ^78^, calculating the decreasing percentage in wound area over time.

---

**Figure S1.**
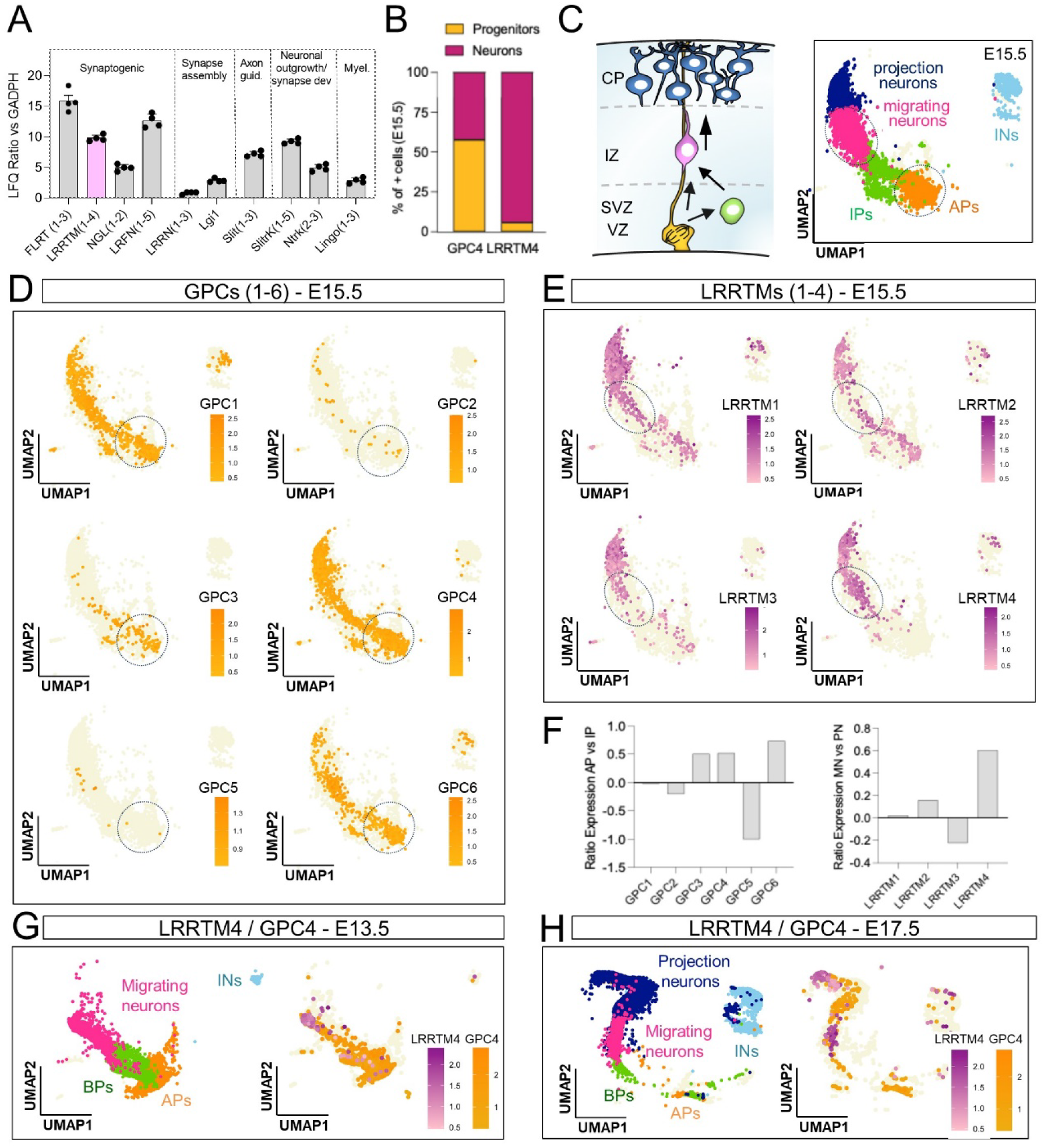
LRRTMs and glypicans expression during cortical development. (A) LRR-containing receptors identified from E15.5 mouse cortices by lectin pulldown followed by mass spectrometry using label-free quantification (LFQ). **(B)** Percentage of progenitor cells (yellow) and neurons (magenta) expressing GPC4 and LRRTM4, based on scRNAseq data from E15.5 mouse cortex (GSE153164). **(C)** Schematic representation of the major cell populations identified in the UMAP visualization of scRNA-seq data from E15.5 mouse cortex published in di Bella et al. (2021) (GSE153164) (left). **(D)** Glypican(1-6) mRNA expression per cell mapped on the E15.5 UMAP shown in C. Most glypicans are enriched in the AP cluster (dashed line). **(E)** Similar to D, but for LRRTM(1-4) members. The majority of LRRTM4-expressing cells localize to the migrating neuron cluster (dashed outline). **(F)** Ratio of glypican (GPC1–6) expression in intermediate progenitors relative to apical progenitors (left), and ratio of LRRTM (LRRTM1–4) expression in migrating neurons relative to projection neurons (right). **(G and H)** UMAP visualization of scRNA-seq data from E13.5 (G) and E17.5 (H) mouse cortex (di Bella et al., 2021, GSE153164). Major cell clusters are colored according to cell-type annotations from published metadata (left). Combined plot of GPC4 (yellow) and LRRTM4 (magenta) is shown on the right.

**Figure S2.**
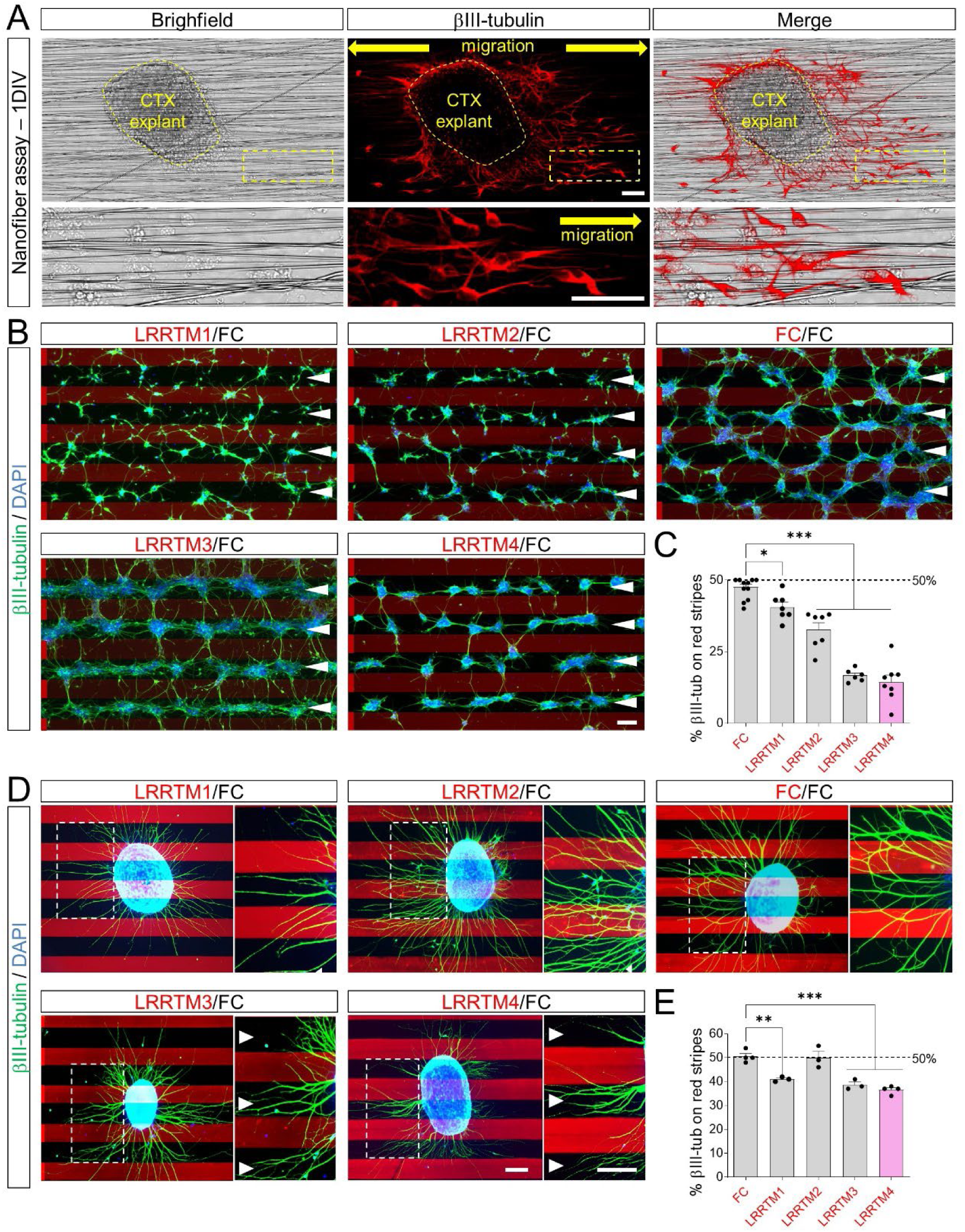
LRRTMs induce repulsion to cortical neurons. **(A)** Cortical explants cultured on nanofibers and stained for βIII-tubulin (red) after 1 day in vitro (DIV). The perimeter of the explant is delineated by a yellow dashed line. Insets (yellow rectangles) highlight migrating neurons extending along the nanofibers. **(B)** E15.5 dissociated cortical neurons were grown on alternate stripes containing FC (black) and LRRTM(1-4) (red). Neurons were stained with anti-β-III-tubulin (green) and nuclei (DAPI, blue). **(C)** The percentage of bIII-tubulin+ pixels on red stripes was quantified. n > 3 different experiments. ∗p < 0.05, ∗∗∗p < 0.001, one-way ANOVA test with Tukey’s post hoc analysis. **(D)** Similar to B, but cortical explants were grown on alternate stripes. Axons were visualized with anti-β-III-tubulin (green). **(E)** The percentage of bIII-tubulin+ pixels on red stripes was quantified. n > 3 different experiments. ∗∗p < 0.01, ∗∗∗p < 0.001, one-way ANOVA test with Tukey’s post hoc analysis. Scale bars represent 50 um (A), 100um (B,D).

**Figure S3.**
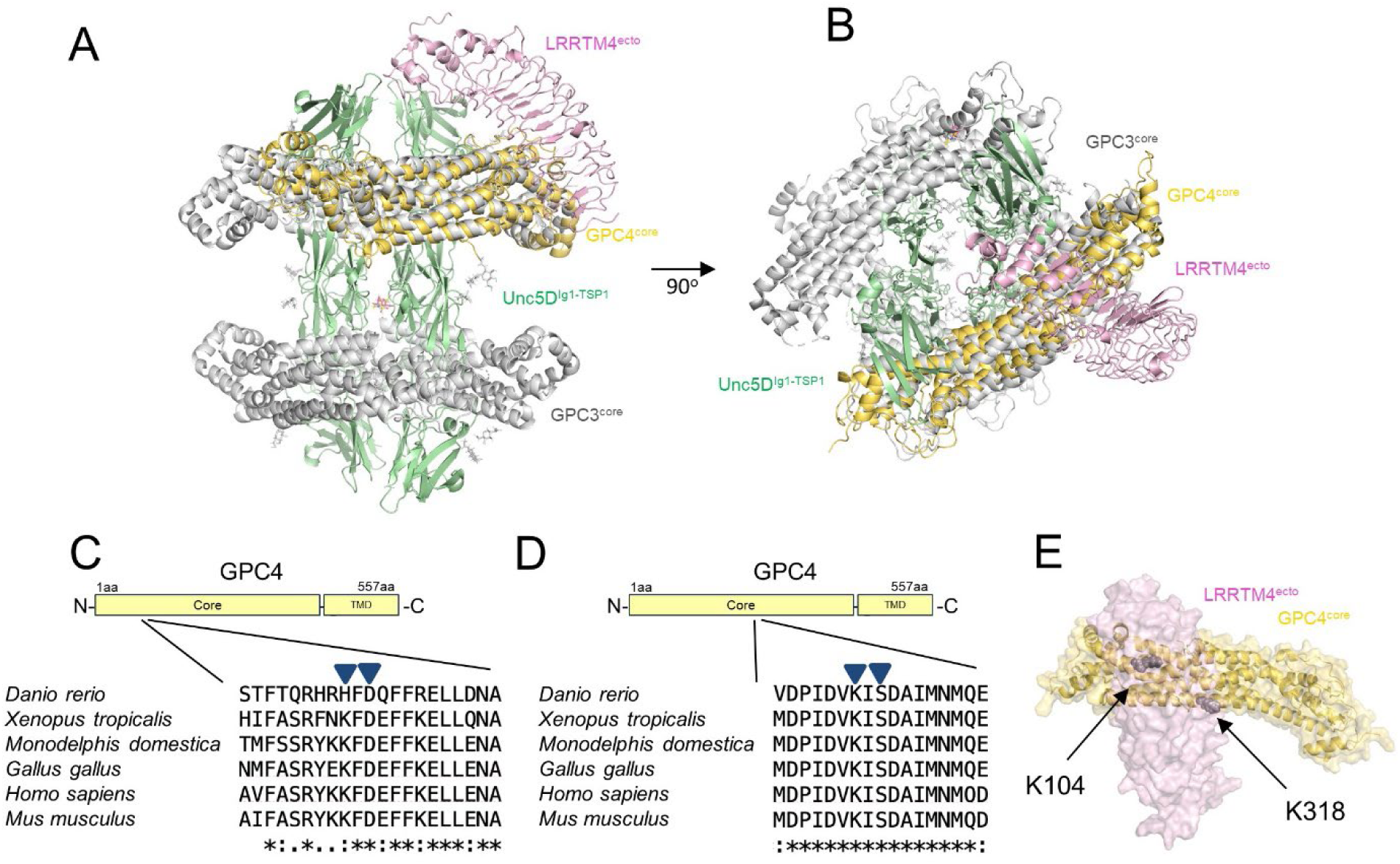
**(A).** Structural superposition of GPC4 (yellow), as predicted to bind to LRRTM4 (pink), and one copy of GPC3 (grey) as found in complex with Unc5D (light green). **(B)** As panel A, but rotated by 90 degrees. **(C)** A sequence alignment highlights the position of K104, which was mutated in the GPC4 mutant K104N+D106T. **(D)** As panel C, but showing the position of K318. **(E)** The structural model shows where K104 and K318 (depicted as grey spheres) are located within the predicted GPC4-LRRTM4 interface.

**Figure S4.**
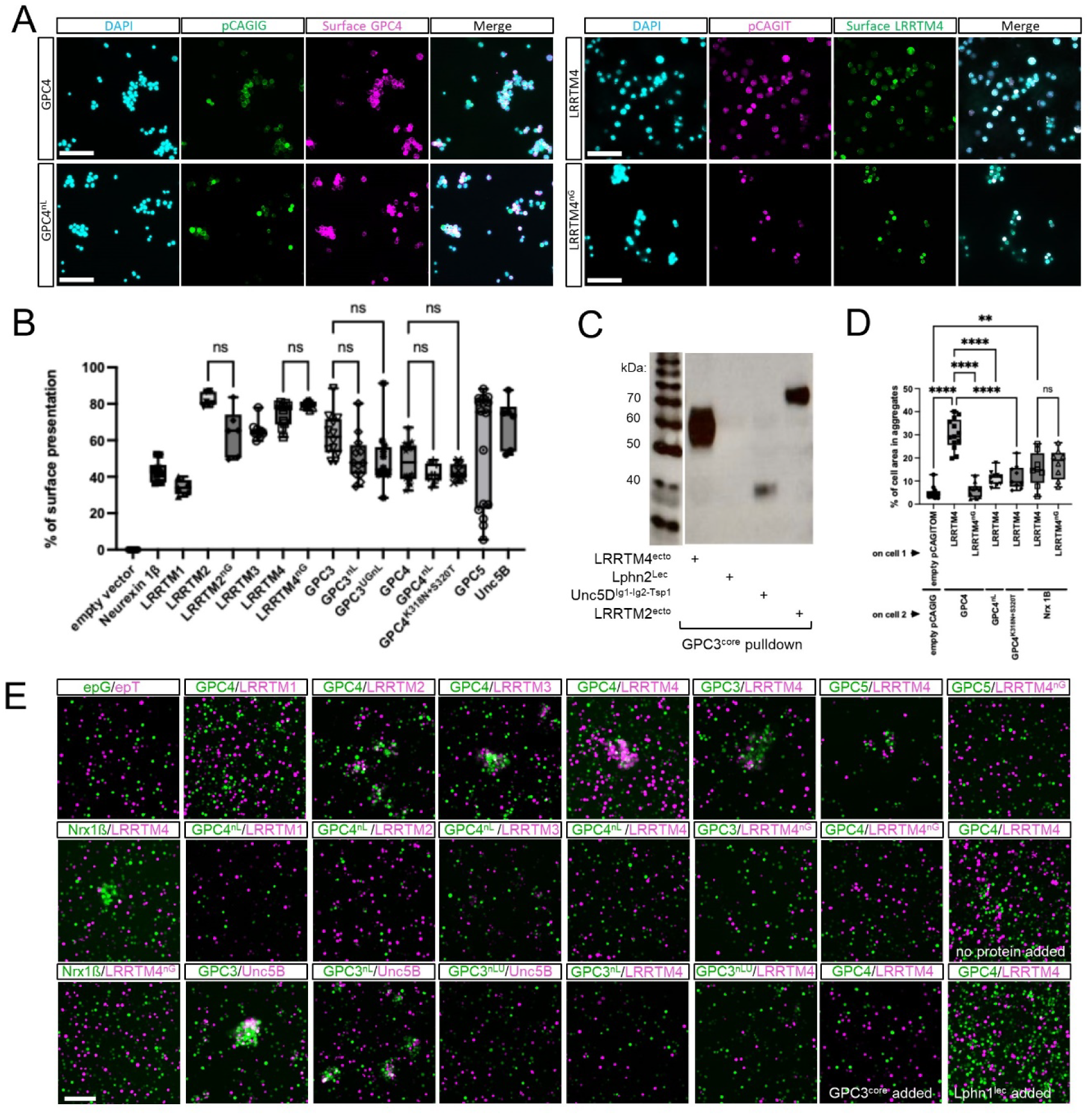
Validation of the GPC-LRRTM interaction and of the structure-based mutants. **(A)**: Representative images of a cell surface staining checking the surface presentation of all the constructs used in Figure 4 for cell aggregation assays. **(B):** Quantification of the cell surface staining experiment in A. **(C):** Pull-down experiment showing that GPC3 interacts with both LRRTM2 and LRRTM4 in solution. **(D)**: Cell-based aggregation assay showing that both GPC4 mutants: K104T+D106T and K318N+S320T impede LRRTM-GPC interaction. (E): Representative images of the cell-based aggregation experiments. n.s. = not significant. ** p<0.01. ****p < 0.0001. One-way ANOVA test with Tukey’s post hoc analysis (B,D). Box and whiskers plots (B,D) are defined as minimum data point to maximum data point, centre is median, percentiles are 25, 50, 75. Scale bars (A,E) represent 100μm.

**Figure S5.**
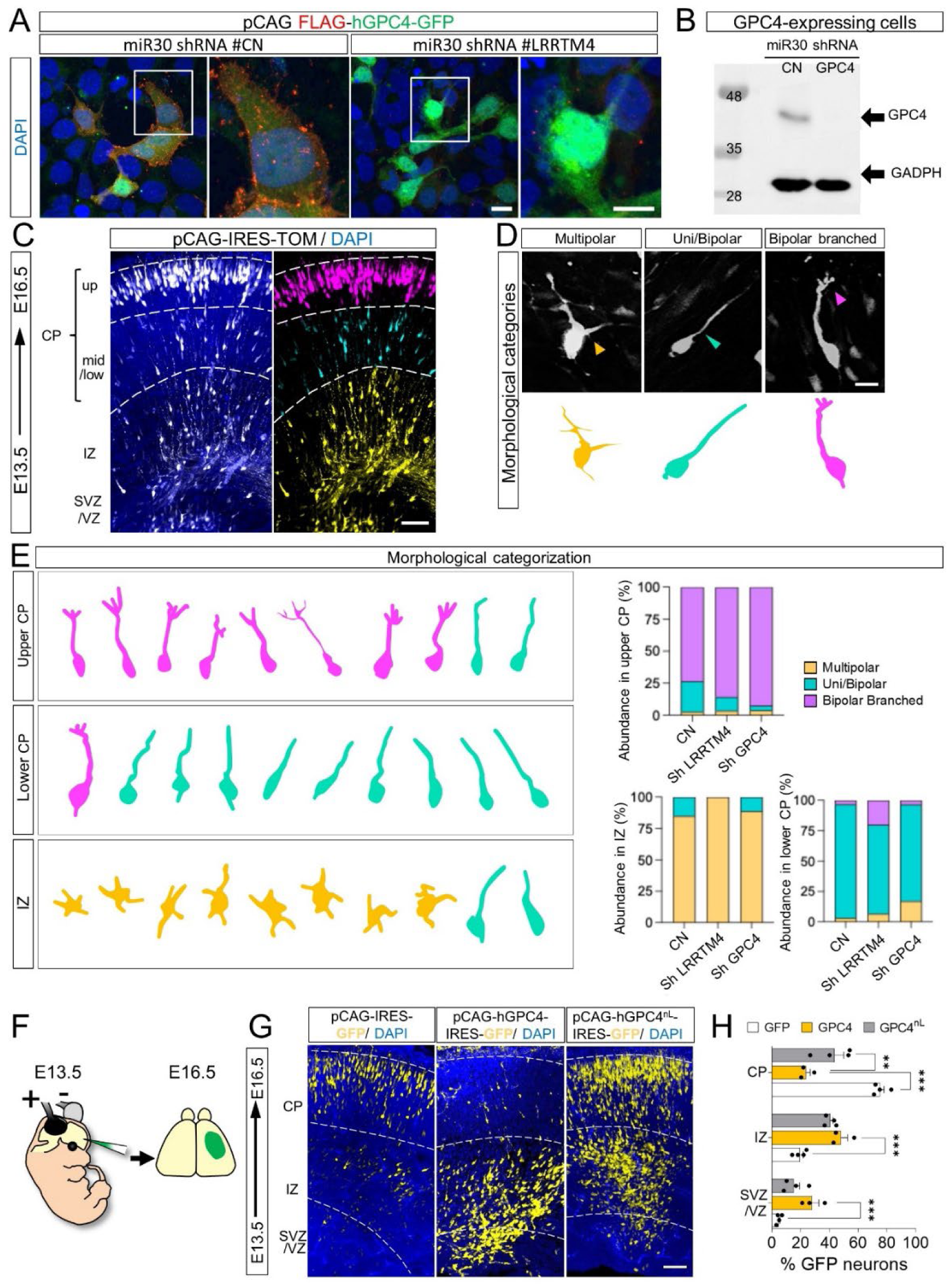
Validation of GPC4 shRNA construct and neuron morphology. **(A)** HeLa cells were transfected with pCAG expressing FLAG-tagged GPC4 (red) together with pCAG-miR30 containing a control shRNA or an shRNA targeting murine GPC4. Cells were cultured for 2 DIV, and immunostained for surface GPC4 (FLAG tag, red). A representative cell for each condition are shown, with magnified views on the right. **(B)** Anti-FLAG western blot showing expression levels of FLAG-tagged GPC4 in the same cells transfected in A. GADPH was used as a loading control. **(C)** Electroporated neurons in the upper CP (magenta), mid-lower CP (cyan) and IZ (yellow) are colored according to the highest abundance of each morphological category in each bin. Nuclear staining with DAPI is shown in blue. **(D)** We categorized neurons overexpressing nanobodies or GFP into multipolar, uni/bipolar, or bipolar branched phenotypes (example images). **(E)** Electroporated neurons in the upper CP (magenta), mid-lower CP (cyan) and IZ (yellow) are colored according to the highest abundance of each morphological category in each bin. Nuclear staining with DAPI is shown in blue. Abundance of each category of neurons in the upper/mid-lower CP and in the IZ of electroporated brains is shown on the right. **(F)** Scheme of the in utero electroporation used in panel G. **(G)** Coronal sections of E16.5 cortex after IUE using empty vector (pCAGIG, control), GPC4 and GPC4nL. GFP-positive cells were quantified for each bin. **(H)** Quantification of data shown in (E). n = 4 GFP, n = 4 GPC4, and n = 4 GPC4nL electroporated brains. ∗p < 0.05, ∗∗p < 0.01, ∗∗∗p < 0.001, one-way ANOVA test with Tukey’s post hoc analysis. Scale bar 10um (A,D), 100um (C,G).

**Figure S6.**
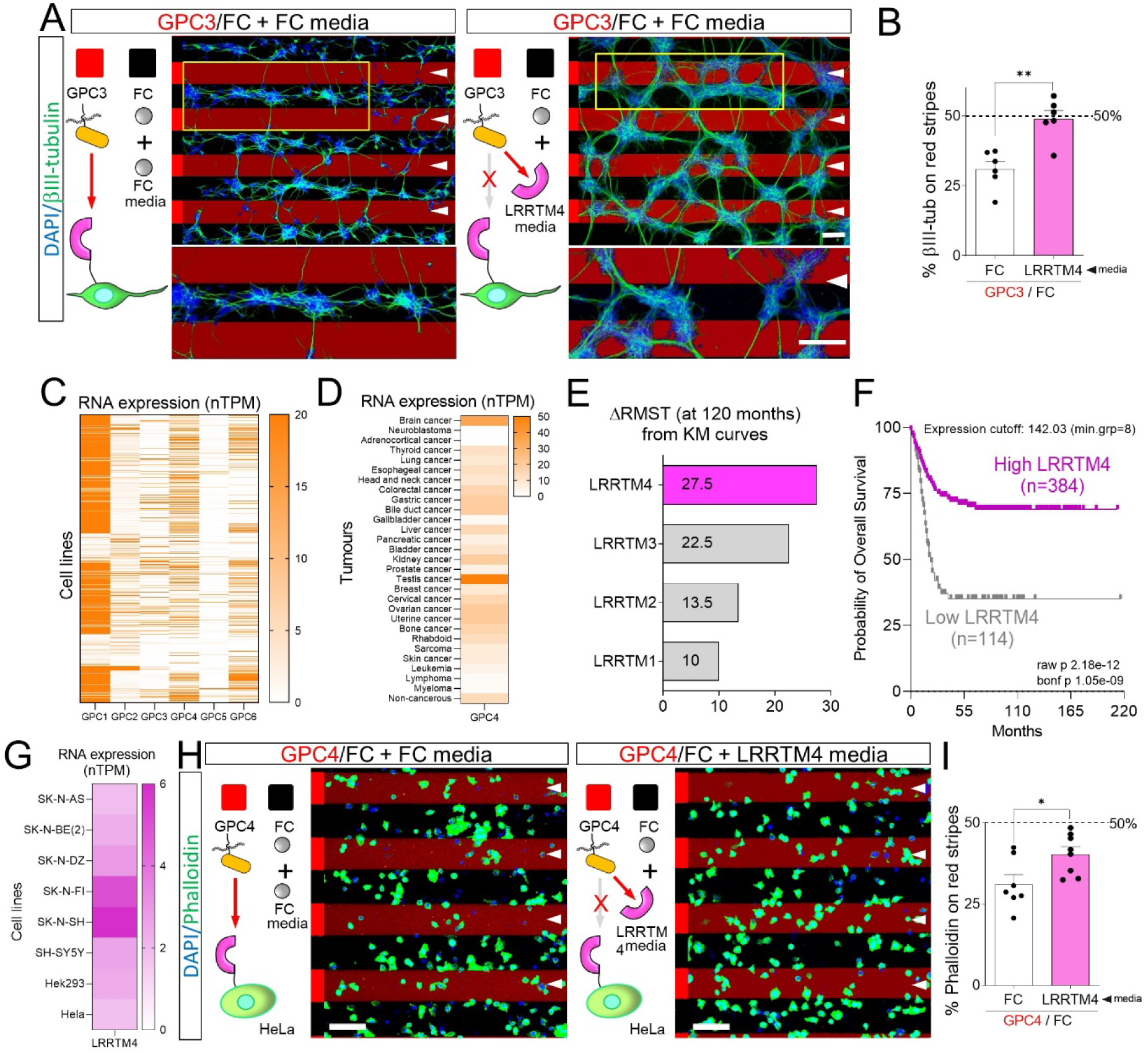
LRRTM4 contributes to Glypican-dependent repulsion in cancer cells. **(A)** E15.5 dissociated cortical neurons grown on alternate stripes containing FC (black) and GPC3 proteins (red) in the presence of soluble FC or LRRTM4 ectodomain. Neurons were stained with anti-β-III-tubulin (green) and DAPI (nuclei, white). Red stripes are indicated by white arrowheads. Yellow rectangle is magnified on the bottom. **(B)** Quantification of the percentage of β-III-tubulin pixels on red stripes. n > 3 independent experiments. ∗∗p < 0.01, Student’s t test. **(C)** Glypican (1-6) RNA expression in 997 human cancer cell lines, shown as normalized transcripts per million (nTPM). **(D)** As in panel C, but for 32 tumor types from the Cancer Genome Atlas. **(E)** Difference in restricted mean survival time (ΔRMST) between high- and low-expression groups for LRRTM1-4, evaluated at 120 months. Positive values denote improved survival associated with higher expression. **(F)** Kaplan-Meier survival analysis of LRRTM4 expression in neuroblastoma. Patients were stratified into high (n = 384) and low (n = 114) LRRTM4 expression groups. **(G)** LRRTM4 RNA expression in several cell lines. **(H)** Dissociated HeLa cells grown on alternate stripes containing FC (black) and GPC4 proteins (red) in the presence of soluble FC or LRRTM4 ectodomain. Cells were stained with phalloidin (green) and DAPI (nuclei, blue). Red stripes are indicated by white arrowheads. **(I)** Quantification of the percentage of phalloidin pixels on red stripes. n > 3 independent experiments. ∗p < 0.05, Student’s t test. Scale bar 100um (A,G).

